# Hippocampal astrocytic sequences emerge during learning and memory

**DOI:** 10.1101/2024.09.06.611660

**Authors:** Ryan A. Senne, Rebecca L. Suthard, Rui Cao, Siria Coello, Amy H. Monasterio, Evan A. Ruesch, Michelle D. Buzharsky, Marc W. Howard, Steve Ramirez

**Affiliations:** Graduate Program for Neuroscience, Boston University, Boston, MA 02215; Department of Biomedical Engineering, Boston University, Boston, MA 02215; Department of Psychological and Brain Sciences, The Center for Systems Neuroscience, Neurophotonics Center, and Photonics Center, Boston University, Boston, MA 02215; Department of Brain and Cognitive Sciences, Massachusetts Institute of Technology, Cambridge, MA 02139; Department of Psychological and Brain Sciences, University of Utah, Salt Lake City, UT 84112; Department of Psychology and Neurosciences, University of Tennessee, Knoxville, TN 37961

## Abstract

The dorsal hippocampus is a heterogeneous structure with numerous cell types involved in generating and maintaining detailed representations of space and time. Prior work has established that pyramidal cells contribute to these crucial aspects of associative memory. For example, hippocampal “time cells” encode temporal information through sequential activity. However, the roles of non-neuronal cell types are less often explored. In this study, we investigated dorsal hippocampal CA1 astrocytes using one-photon calcium imaging in freely moving animals during a contextual fear conditioning paradigm, enabling tracking of a population of glial cells across days. In response to foot shock, astrocytes generated robust calcium-event sequences with a time-compressed structure akin to canonical hippocampal time cells. Upon re-exposure to the conditioned context, we again detected astrocytic sequences -- in the absence of shock -- displaying similar time-compressed structure. Importantly, this phenomenon was not observed in a context different from the initial fear conditioning chamber. Lastly, spontaneous astrocytic activity in the initial conditioning context was associated with behavioral state transitions (e.g. freezing), but this was not observed in a novel context. Taken together, these results suggest a direct role for astrocytes in generating hippocampal representations of associative memory, and future work may continue to study how diverse cell types process temporal features of associative memory.

## Introduction.

To create associative memories with high granularity, the hippocampus encodes information by indexing the spatial and temporal aspects of an experience. Within the hippocampus, pyramidal cells function as place and time cells, developing spatially-tuned receptive fields within an environment and tiling temporal epochs through sequential activity, respectively^1–6^. Glial cells, on the other hand, have been largely ignored in this realm of spatiotemporal processing until recently. Recent findings have shown that astrocytes interface with neurons in a synergistic and complementary manner to represent spatial information in dorsal CA1 of the hippocampus (dCA1) and may encode reward location via similar spatial mechanisms^7,8^. Further, computational modeling has proposed that astrocytic neuromodulation increases the number of active place fields and refines neuronal encoding structure via enhanced theta oscillations and plasticity changes^9^. Together, these findings point to astrocytic involvement in spatiotemporal processing during memory formation and retrieval, and how it may relate to neuronal activity within these circuits.

It remains unclear whether astrocytic activity is temporally organized in a way that is directly relevant to hippocampus-dependent learning and recall. This question is particularly salient for contextual fear conditioning, in which the dorsal hippocampus is essential for linking a context with an aversive event and subsequently retrieving that association. Recent studies have implicated astrocytes in learning and memory through perturbation of astrocytic signaling and gliotransmission, and have suggested causal roles for astrocyte-neuronal ensemble interfaces in memory stability and recall^10–14^. However, despite this evidence, comparatively little work has directly measured astrocyte population dynamics at single-cell resolution in dCA1 during fear learning and recall. Consequently, it is still unknown whether astrocytes exhibit structured activity around the onset of an aversive event, whether such structure is stable across days, whether it re-emerges during memory retrieval, and whether extant temporal organization relates to canonical hippocampal coding motifs and fear-related behavior (e.g., freezing). Only recently have calcium imaging–based methods enabled these questions to be addressed directly in freely-behaving animals.

Here, we hypothesized that single-cell astrocyte dynamics in dCA1 carry organized signatures of fear-related events and memory retrieval. To test this, we used longitudinal one-photon calcium imaging to record activity from thousands of individual dCA1 astrocytes across learning and memory in a contextual fear conditioning (FC) paradigm, tracking the same cells across conditioning and subsequent contextual recall. This approach allowed us to ask if astrocyte activity is time-locked to aversive foot shocks, if astrocytic population dynamics exhibit motifs reminiscent of coding observed in neuronal ensembles, and if these dynamics reappear during retrieval in a manner that relates to ongoing behavior.

Using this framework, we found that dCA1 astrocytes are shock-responsive at the single-cell level and that aversive foot shock onset triggers structured astrocyte population sequences during conditioning. Surprisingly, these sequences displayed features consistent with temporally organized ensemble activity described for canonical hippocampal time cells. During contextual recall, related sequence structure reappeared and covaried with freezing behavior, suggesting that astrocytic temporal dynamics are associated with behaviorally relevant aspects of memory retrieval. However, such activity may also reflect broader state transitions that accompany memory expression, like arousal. Together, these findings support a role for astrocytes in temporally structured processing within hippocampal circuits and motivate further study of how astrocytic and neuronal dynamics interact to shape and express memory.

## Results

### One-photon imaging of dorsal hippocampal astrocytes across contextual fear learning

To examine astrocytic activity during contextual fear conditioning (CFC), we performed one-photon (1p) calcium imaging of dorsal CA1 (dCA1) astrocytes in freely-moving mice (**Fig. 1a)**. Wild type mice were injected with an astrocyte-specific genetically-encoded calcium indicator (AAV5-GfaABC1D-cyto-GCaMP6f), and a gradient-index (GRIN) lens was implanted above the pyramidal cell layer to enable longitudinal imaging of astrocytic calcium activity (**Fig. 1a-c**). To verify astrocyte-specific expression, we performed immunohistochemical co-staining for glial fibrillary acidic protein (GFAP) or NeuN with GCaMP6f beneath the GRIN lens. Of the 1326 GCaMP6f-expressing cells, only 3 were NeuN-positive neurons, yielding an off-target rate of 0.23%. Conversely, 1204 of 1206 GCaMP6f-expressing cells were GFAP-positive astrocytes (99.8% specificity; **Extended Data Fig. 1a-b**), confirming highly selective astrocytic expression.

**Fig. 1:**
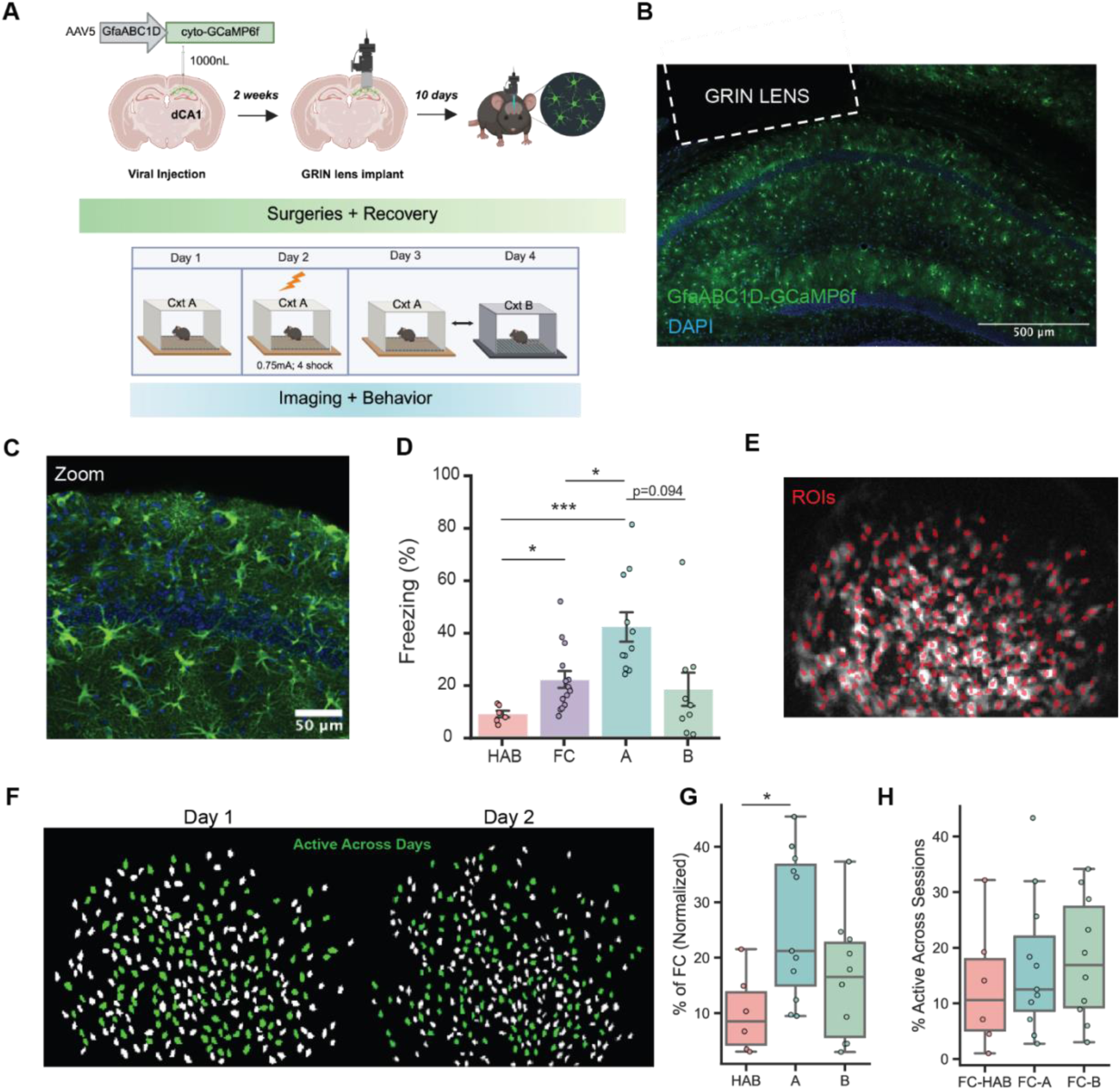
One-photon imaging of dorsal hippocampus astrocytes during contextual fear learning and recall. (A) Experimental timeline. Wild-type C57BL/6J mice expressing GCaMP6f selectively in astrocytes in dorsal CA1 were implanted with a GRIN lens and imaged using one-photon microscopy during habituation, contextual fear conditioning (FC), and recall. During FC, mice received four foot shocks (0.75 mA, 2 s) at defined time points. Recall was tested in the conditioned context (Context A) or a novel context (Context B) in a counterbalanced design. (B) Representative confocal image of the dorsal CA1 pyramidal cell layer showing astrocytic GCaMP6f expression (green) and DAPI-labeled nuclei (blue), with approximate GRIN lens placement indicated. Scale bar, 500 µm. (C) Higher-magnification image illustrating astrocyte morphology and GCaMP6f expression. Scale bar, 50 µm. (D) Freezing behavior across habituation, FC, and recall sessions. Freezing was increased during FC and recall in Context A compared to habituation, but not in Context B (linear mixed effects model with Bonferroni correction). (E) Representative regions-of-interest (ROIs) showing astrocytes active during FC from a single mouse. ROIs were manually curated (*see Methods*). (F) Example CellReg output illustrating longitudinal registration of astrocytes across sessions; green ROIs indicate cells detected in both sessions. (G) Proportion of astrocytes active during each session, normalized to FC. A greater proportion of astrocytes were active during recall in Context A compared to habituation. (One-way ANOVA (logit) with Bonferroni correction). (H) Proportion of astrocytes detected across session pairs. No significant differences were observed between session pairs. (One-way ANOVA (logit)). Error bars indicate mean ± SEM. Significance levels: *p ≤ 0.05, **p ≤ 0.01, ***p ≤ 0.001, ****p ≤ 0.0001. HAB (n = 7), FC (n = 14), Context A (n = 11), Context B (n = 10). See Supplemental Table 1 for statistical details.

Behavioral testing occurred over four days (**Fig. 1a**). On Day 1, mice underwent habituation (HAB; i.e., pre-exposure to Context A (Cxt A)) in the absence of aversive stimuli. On Day 2, all mice underwent FC in Cxt A to acquire the association between the conditioned stimulus (context; CS) and unconditioned stimulus (foot shock; US) which were paired at the 120, 180, 240 and 300s time points during a 330s session (foot shocks; 2s duration; 0.75mA intensity). On Days 3 and 4, mice were either placed back into Cxt A (recall) or into a novel Context B (Cxt B) to control for context-specificity and time-dependent effects. These days were counterbalanced where applicable.

Freezing behavior differed significantly across sessions (**Fig. 1d**). A linear mixed effects model with animal as a random effect revealed a significant main effect of context, with freezing significantly higher in Cxt A and FC relative to the HAB session (Cxt A: β = 35.61, p<0.0001; FC: β = 15.85, p = 0.037), but not in Cxt B (β = 11.38, p = 0.150). *Post hoc* multiple comparisons confirmed higher freezing in Cxt A vs. HAB (p=0.008) and FC vs HAB (p=0.013), with no difference between Cxt B vs. HAB (p=1.000). Freezing was also significantly higher in Cxt A compared to FC (p=0.043), and importantly, there was evidence of some degree of generalization across Cxt A and Cxt B (p=0.09) (Bonferroni) (**Fig. 1d**). Importantly, freezing level was not significantly impacted by sex or counterbalancing order, but varied by session, increasing during Cxt A (β = 37.65, p<0.001) and FC (β = 17.84, p = 0.019), with no changes in Cxt B (β = 12.04, p = 0.124) (linear mixed effect model; **Extended Data Fig. 2a**).

To assess astrocytic activity across learning, we longitudinally tracked regions-of-interest (ROIs) from each imaging session. Astrocytes were reliably matched across days using a combination of manual ROI identification (*see Methods*) and CellReg, which aligns cellular footprints across days^15^. This approach enabled classification of astrocytes that were active exclusively on Day 1 or Day 2, as well as those that were active across both sessions. A representative field-of-view illustrates all astrocytes detected during FC using manual ROI selection (**Fig. 1e**), and a corresponding CellReg output highlights those astrocytes tracked across sessions (green ROIs; **Fig. 1f**). When normalized to the FC session, the proportion of active astrocytes differed across HAB, Cxt A and Cxt B (One-way ANOVA (logit-transformed data): F(2,24) = 6.68, p = 0.024), driven by reduced activity in HAB relative to Cxt A (p=0.018, Bonferroni), with no other pairwise differences (**Fig. 1g**). Further, longitudinal tracking of astrocytes enabled quantification of the fraction of cells active across session pairs (HAB-FC, FC-Cxt A, FC-Cxt B). The proportion of astrocytes active across days did not differ significantly (One-way ANOVA (logit-transformed data): F(2,24) = 1.54, p=0.512; **Fig. 1h).** Importantly, astrocyte recruitment (% active across days) did not differ by counterbalancing order or session pair, and no interaction was observed (two-way ANOVA, interaction: F(2,21)=1.62, p=0.220; session pair: F(2,21)=0.72, p=0.500; order: F(1,21)=0.08, p=0.770; **Extended Data Fig. 2b, right**). In contrast, a significant main effect of sex was detected, with no interaction between sex and session pair (two-way ANOVA, sex: F(1,21)=4.84, p=0.039; interaction: F(2,21)=0.12, p=0.890; **Extended Data Fig. 2b, left)**. Together, these results demonstrate reliable longitudinal tracking of astrocytes during one-photon imaging and indicate that comparable fractions of astrocytes are active between FC and subsequent recall sessions, independent of context.

### Evidence of astrocytic temporal sequences after foot shock onset

To understand the nature of astrocytic single-cell activity during FC, we first visualized individual traces to visualize broad patterns across the session (**Fig. 2a**). Astrocytes exhibited robust responses to foot shock, but with heterogeneous profiles in their individual cell calcium physiology (**Fig. 2a**). Qualitatively, individual astrocytes displayed unique kinetics and differences in the number of shocks they responded to within a session, although nearly all cells (98%) responded to at least one shock (**Fig. 2a-b**). At the population level, we observed that astrocytic calcium activity was significantly elevated after the onset of each foot shock, peaking around 5 seconds post-stimulus (**Fig. 2b**; peri-event analysis). This finding is in line with previous work from our lab and others showing astrocytic responses to aversive stimuli in the hippocampus, cortex, and amygdala^16–18^.

**Fig. 2:**
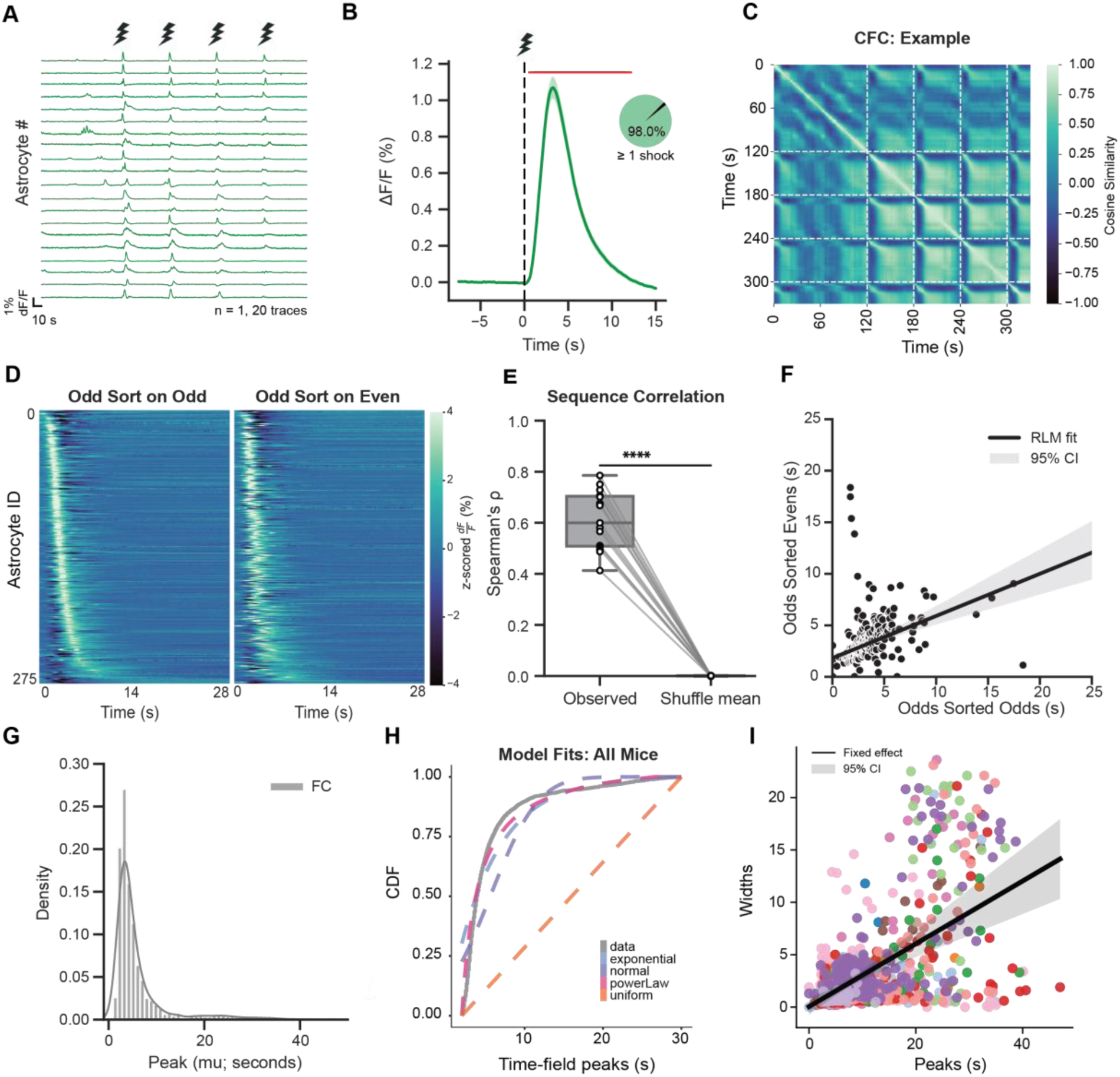
Astrocytic temporal sequences emerge following foot shock onset. (A) Example astrocytic calcium traces (n = 20 cells; n = 1 mouse) during contextual fear conditioning, showing robust responses to foot shock. Fluorescence is expressed as ΔF/F (%). Scale bar, 10 s. (B) Peri-event averaged astrocytic calcium activity aligned to foot shock onset (time = 0; dashed line). Traces were averaged within astrocytes and then across animals (n = 14 mice). Shaded regions indicate 95% confidence intervals; red bars denote periods of significant deviation from baseline. *Inset*, proportion of astrocytes responsive to at least one foot shock. (C) Example cosine similarity matrix showing pairwise similarity of astrocytic calcium activity from a single mouse, revealing block-like temporal structure emerging after the first foot shock. Dashed lines indicate shock times. (D) Heatmap of astrocytic calcium activity from a single mouse aligned to foot shock onset. Astrocytes were sorted by peak response time during odd-numbered shocks and applied to even-numbered shocks to assess sequence stability. Stable temporal organization was observed across animals (n = 14). (E) Spearman rank correlations between peak response times derived from odd trials and cross-validated peak times derived from even trials. Each point represents one animal (n = 14). Correlations were significantly higher than a shuffle null distribution (Paired t-test). (F) Robust linear regression from an example animal illustrating preserved temporal ordering of astrocytic peak times across cross-validation. (G) Distribution of predicted astrocytic peak times (µ; seconds) across the population (n = 14 mice), revealing a skewed temporal organization following foot shock onset. (H) Empirical cumulative distribution function (ECDF) of predicted peak times with fitted theoretical distributions. Model comparisons favored a Power-law distribution (pink; WAIC=12008.7) over exponential (blue; WAIC=14423.3), normal (purple; WAIC=15832.2), and uniform (orange; WAIC= 17627.4), consistent with temporally compressed structure. (I) Relationship between astrocytic time field width (σ) and peak response time (µ) for individual astrocytes across all animals. Width increased with peak location, consistent with temporal scaling. Line indicates the fixed effects best fit from a linear mixed model with 95% CI (n = 14; cells = 2707). Colored dots represent observations from a particular animal. Significance levels: *p ≤ 0.05, **p ≤ 0.01, ***p ≤ 0.001, ****p ≤ 0.0001. See Supplemental Table 1 for statistical details.

### Robust astrocytic sequences emerge and tile the interval after foot shock for up to tens of seconds

To examine the basic structure of population activity, we computed the cosine similarity of the astrocyte calcium population vector over time **(Fig. 2c, Extended Data Fig. 3).** Broadly, we noted a set of recurring motifs present in the cosine similarity plots; at the timing of the initial foot shock, a block-like structure emerged and repeated across subsequent shock presentations and higher off-diagonal similarity between timings of foot shock. Such off-diagonal structure is characteristic of populations exhibiting sequential activity, suggesting that astrocytes may engage in sequence-like dynamics in response to foot shock during learning. However, it is worth noting that these features did have variability across animals, necessitating further analysis **(Fig. 2c, Extended Data Fig. 3)**.

Before analyzing astrocytic responses to shock, we asked whether locomotion-related signals were represented at the population level, given prior evidence that movement accounts for a substantial fraction of astrocyte calcium activity^19,20^. To minimize potential contamination from fear-related signals, we restricted this analysis to 120 seconds preceding the onset of the first foot shock **(Extended Data Fig. 4a)**. We then used principal component analysis (PCA) to assess how velocity information was embedded in population activity **(Extended Data Fig. 4b)**. Across animals (n = 6), velocity-related structure was largely absent; only a single animal showed clear evidence of movement encoding. In this animal, velocity was reflected in the subspace score (*see Methods*) and could be decoded from the top ten principal components at levels exceeding chance **(Extended Data Fig. 4c–d)**. Consistent with these population-level results, this same animal was also the only case in which the mean single-cell correlation with velocity was reliably greater than zero **(Extended Data Fig. 4e)**. Finally, velocity-related variance in this animal was concentrated primarily in the first principal component, which correlated with velocity at approximately r ≈ 0.55 **(Extended Data Fig. 4f)**. These results provide evidence that our signals are not primarily driven by locomotion-specific information.

Given that locomotion was not generally a reliable indicator of astrocytic activity, we next plotted the same activity for the odd-numbered shocks and ordered the traces by each astrocyte’s peak calcium response time during Shocks #2 and #4 (even-numbered shocks) **(Fig. 2d, right; Extended Data Fig. 5)**. To quantify the stability of these potential ‘sequences’, we computed Spearman correlation coefficients between the “true” peak times (odd trials ordered using odd-trial peak times) and the cross-validated peak times (odd trials ordered using even-trial peak times) for each animal. To test whether these correlations exceeded chance at the population level, we generated an empirical null distribution for each animal by randomly permuting the cell identities across comparison groups 10,000 times and recomputing rho (ρ) on each shuffle. We then compared the observed ρ values against these animal-specific null means using a paired t-test, which indicated that correlations were reliably above chance **(Fig. 2e**; t = 20.53, p<0.0001). We also note that all animals were significantly above chance compared to their observed statistics against their own respective null distributions (**see Supplemental Table 1**). Next, we fit a linear regression model for all animals to determine if these true peak times and the cross-validated peak times were related. We opted for a robust Huber regression model, given the presence of clear heteroscedasticity in the data, and found a significant relationship between these variables (β =0.41, p<0.0001; **Fig. 2f; Extended Data Fig. 5)**. To further evaluate how these relationships changed as a function of stimulus presentation, we performed the same analysis on the early (Shock #1 and #2) against the late (Shock #3 and #4) shocks. We found that the temporal order of astrocytic responses is stable across the course of conditioning (paired t-test; t=18.79, p<0.0001; **Extended Data Fig. 6).**

Using a hierarchical Bayesian model, we fit each astrocyte’s post–shock response with a time-field function parameterized by a peak time (µ) and a receptive-field width (σ) (*see Methods for model details*); similar to approaches used to characterize hippocampal time cells^21^. To summarize the distribution of inferred peak times, we first visualized µ across all animals during FC (n = 14) and observed a strongly right-skewed distribution following shock onset **(Fig. 2g)**, consistent with a temporally compressed representation. Sorting the average astrocytic responses by µ revealed robust post-shock sequences in every mouse, extending for tens of seconds (median peak = 4.1 s; IQR = 3.0–6.4 s; 90th percentile = 12.1 s; **Extended Data Fig. 7)**.

To formally test whether the population peak-time distributions exhibited a time-cell-like structure rather than reflecting noisy peak estimates, we fit the empirical peak times distributions with several candidate models (Power-law, Exponential, Uniform, and Gaussian), excluding cells peaking during shock delivery (<2 s; 1.7% of cells). Model performance was evaluated using the Watanabe–Akaike Information Criterion (WAIC), where lower values indicate a better fit, and visualized via the corresponding cumulative distribution functions **(Fig. 2h)**^22^. The Uniform model provided the poorest fit (WAIC = 17627.4), arguing against the interpretation that the apparent sequences are homogeneously scattered across the session length. This was followed by the Gaussian model (WAIC=15832.2) which argues against an interpretation where peak locations are independently sampled around some location. In contrast, models commonly implicated in hippocampal time-cell sequences—particularly the Power-law—fit substantially better (WAIC = 12008.7) **(Fig. 2h)**^21,23^.

Finally, we asked whether later peaks were associated with broader time fields, a hallmark of temporal compression. We fit a random-slope and intercept linear mixed-effects model in which σ was regressed on µ **(Fig. 2i)**. Receptive-field width increased linearly with peak time (slope = 0.30, p<0.001), indicating progressively reduced temporal resolution for later components of the sequence. Together, these results provide quantitative support that post-shock astrocyte sequences are temporally compressed, paralleling core properties reported for hippocampal time-cell representations^6,21,24,25^.

### Model agnostic analysis of astrocyte calcium activity

Having established that astrocyte population activity exhibits reliable sequential organization after shock onset, we next asked whether an orthogonal, model-agnostic approach would yield a consistent description of the heterogeneity observed at the single-cell level. To this end, we applied WaveMAP, an unsupervised dimensionality reduction and clustering framework designed to identify distinct waveform motifs in electrophysiology and calcium imaging data (**Extended Data Fig. 8**)^26,27^. We pooled astrocyte responses across animals during conditioning (extracted peri-shock windows (–7.5 to 15 s), computed each cell’s average shock-evoked trace, and z-scored traces prior to WaveMAP embedding (*see Methods*).

WaveMAP identified six calcium response clusters (**Extended Data Fig. 8a, c-d**). Notably, when cluster-average waveforms were plotted in order of their peak latency, cluster peaks progressed systematically across time, and later-peaking clusters exhibited broader responses (**Extended Data Fig. 8b**). Thus, rather than indicating discrete, context- or animal-specific subpopulations, the clustering largely segmented the response space along two features already suggested by the sequence analyses (e.g., peak location and event width), providing complementary findings of the continuous temporal organization in shock-evoked astrocyte activity. Importantly, clusters were not driven by individual animals, as cells from all animals were represented across clusters (**Extended Data Fig. 8e**).

### Non-parametric analysis reveals putative calcium events in both contexts

We next asked whether temporally structured astrocytic sequences were present during contextual recall. We first visualized representative astrocyte calcium traces during recall in animals returned to the conditioned context (Cxt A) versus those placed in a novel context (Cxt B) **(Fig. 3a)**. Qualitatively, astrocytes in Cxt A exhibited more prominent calcium events than those in Cxt B, consistent with stronger recruitment of population activity upon re-exposure to the conditioned environment. To evaluate population-level structure during recall, we computed cosine similarity matrices as in the conditioning session. Example animals illustrate that both contexts can exhibit off-diagonal structure, suggestive of repeated temporal patterns, but this organization appeared more pronounced in Cxt A than in Cxt B **(Fig. 3b–c; Extended Data Fig. 9-10)**. However, unlike FC, recall sessions lacked a discrete external trigger (e.g., foot shocks), making it difficult to anchor sequence reliability to a known event time and necessitating an unbiased approach to identify candidate sequence occurrences.

**Fig. 3:**
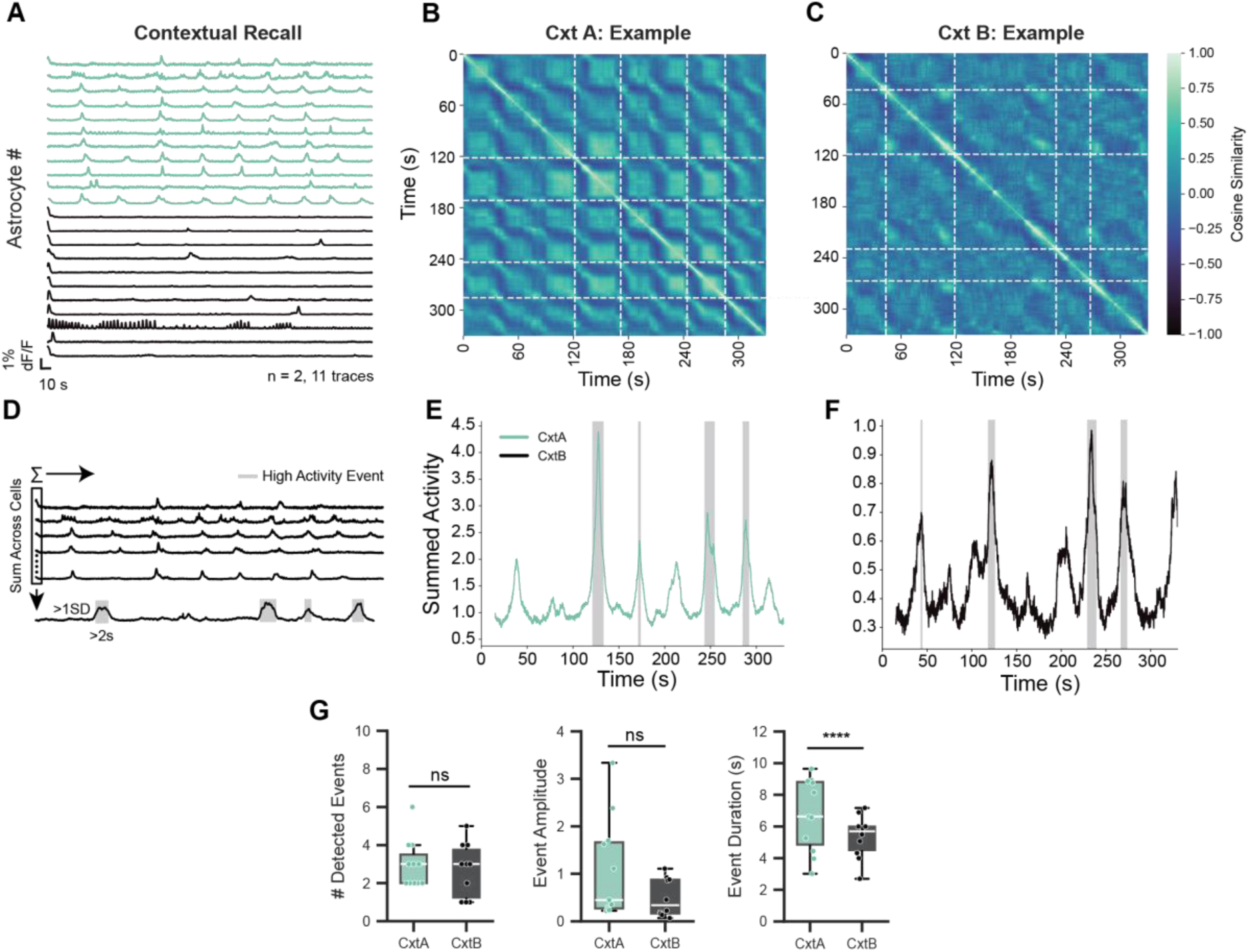
Non-parametric event detection method discovers candidate calcium events in both contexts (A) Example astrocytic calcium traces during recall in Context A (blue; n=20 cells; n=1 mouse) and Context B (black; n=20 cells, n=1 mouse), illustrating distinct activity profiles. Fluorescence is shown as dF/F (%). Scale bar, 10s. (B-C) Example of cosine similarity matrices during recall in Context A (B) and Context B (C), revealing differences in temporal organization. Dashed lines indicate detected high-activity events described in (D). (D) Schematic illustrating spontaneous event detection. Population astrocytic activity was computed by summing activity across cells, and high-activity epochs were detected based on amplitude (>1 SD) and duration (>2 second) thresholds. (E-F) Example population activity traces during recall in Context A (E) and Context B (F), with detected spontaneous calcium events highlighted (grey). Spontaneous high-activity events occurred in both contexts but differed in temporal organization. (G) Number of spontaneous calcium events, event duration, and event amplitude during recall in Context A (n = 11) and Context B (n = 10). There were no significant differences in the detected event number or amplitude between contexts (linear mixed effects model). However, there was a significant effect of event duration, with a shorter duration in Cxt B relative to Cxt A.

Since no external stimulus was present during recall sessions, we next sought to detect candidate spontaneously occurring sequences. Based on our prior FC results, which showed that sequences involve brief periods of strong co-activity, we built an “event detector” based on summed population activity. At each time point, we computed the total activity across cells and labeled points that exceeded 1.0 standard deviations above the mean of the trace. Consecutive points had to last at least 2.0 seconds to be considered a putative “event” **(Fig. 3d)**. To validate this approach, we first applied the detector to FC data and confirmed that all shock-triggered sequences were captured **(Extended Data Fig. 11)**. We then applied the same approach to recall sessions, which revealed high-activity events across all animals and contexts **(Fig. 3e-f** **Extended Data Fig. 12-13)**. Importantly, these detected events represent necessary precursors for sequences, as they reflect periods of elevated population activity, but are not sufficient to establish true sequential structure. Rather, the detector marks candidate epochs of co-activity, but requires further analysis to determine whether they exhibit defining properties of sequences (e.g., temporal compression). With this in mind, we found no significant difference in the number of detected events across contexts (β = −0.21, z = −0.49, p = 0.630), nor in event amplitude (β = −0.587 ± 0.335, z = −1.751, p = 0.080)(**Fig. 3g**, linear mixed-effects model). In contrast, event duration differed significantly, with shorter durations observed in Cxt B relative to Cxt A (β = −1.395 ± 0.178, z = −7.85, p < 0.0001) (**Fig. 3g**, linear mixed-effects model). Together, these findings suggest that contextual differences may not be driven by changes in the frequency or magnitude of population events, but rather by differences in their temporal structure.

### Sequential organization of astrocytic population dynamics is preferential to the conditioned context

We applied the same statistical tests and hierarchical models from FC to evaluate the putative sequences detected during recall. In FC, we found that the peak activity in the sequences induced by foot shocks was correlated across shocks **(Fig. 2d-e).** To determine if this result held during recall, we plotted cross-validated heatmaps of the cellular activity in the same way as during FC (i.e., we plotted the heatmap based on the true sorted peak times (Odd sorted on Odd), and plotted it sorted on the even sequence order (Odd sorted on Even))**(Fig. 4a-b; Extended Data Fig. 14-15)**. Qualitatively, the sorted activity was more structured in Cxt A in comparison to Cxt B. To quantify this, we used the same shuffle control approach used in FC (i.e., randomly permuted cell identities across comparison groups to get an empirical null-mean we could use to compare the observed rho values against). We found that, at the population level, Cxt A animals had significantly greater correlations relative to their shuffle-derived null-values (paired t-test: t=11.57, p<0.0001) whereas Cxt B did not differ from their null distribution (paired t-test: t=1.57, p=0.17) **(Fig. 4c)**. We also found that the null-corrected rho values (subtracting the null-mean from the observed value) of Cxt A was greater than in Cxt B (independent t-test: t=2.60, p=0.032; **Fig. 4c**).

**Fig. 4:**
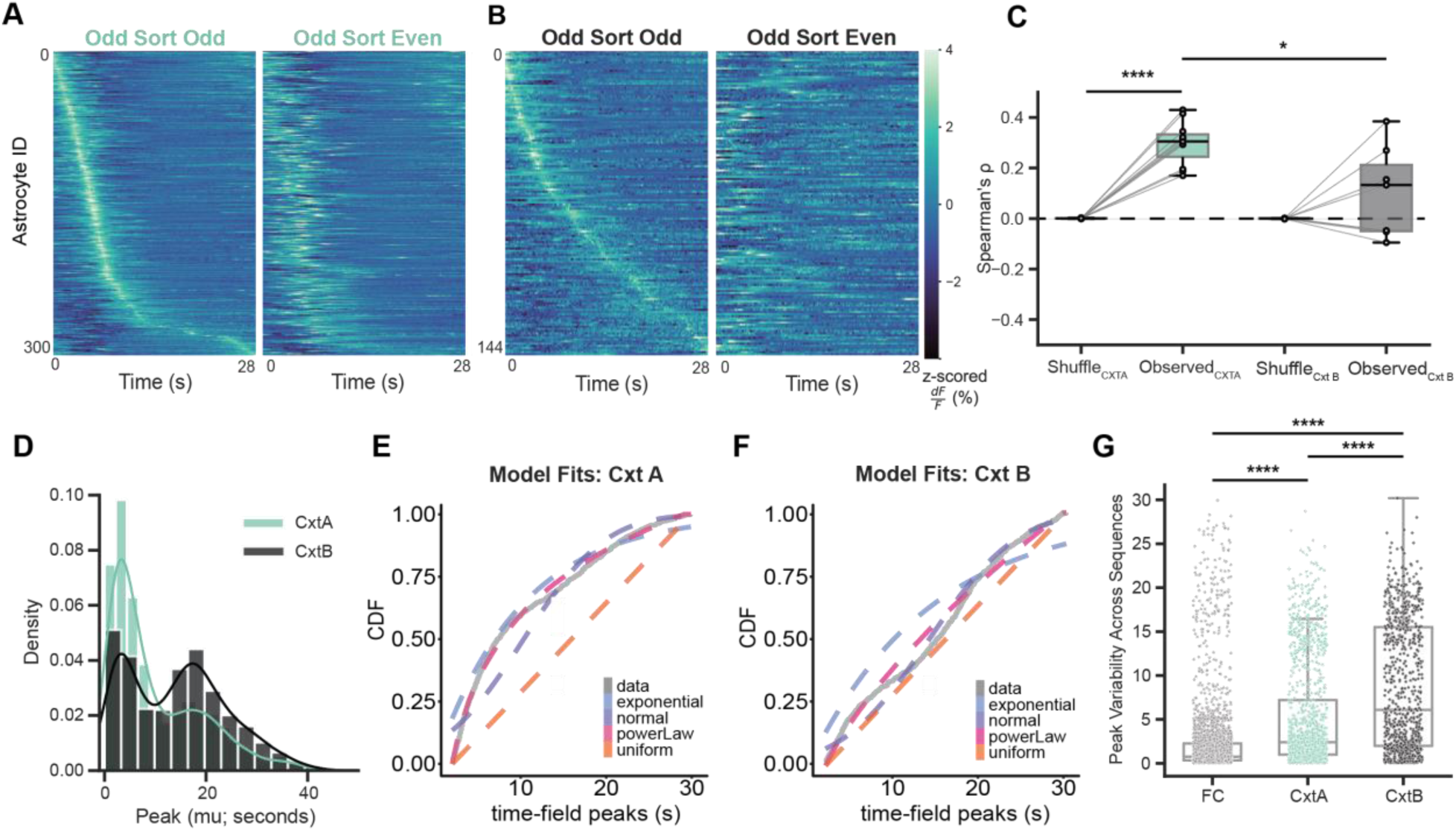
Astrocytes display context-specific sequence structure. (A-B) Example heatmaps of astrocytic calcium activity during spontaneously detected events. Odd-numbered events were sorted by peak timing from odd events (left) or cross-validated using even-numbered events (right) for Context A (A) and Context B (B). Time = 0 is the start time of the detected events on average. (C) Spearman rank correlations between true and cross-validated peak times for Context A and Context B. Correlations exceeded a shuffle-null distribution in Context A but not Context B, indicating context-specific sequence stability (Paired and independent t-tests). Points represent individual animals; grey lines indicate observed-shuffle pairing for each context. Mice with < 2 events were excluded from this analysis. Context A (n = 11) and Context B (n = 7). (D) Population-level distributions of predicted astrocytic peak times (µ; sec) during recall in Context A (blue; n=11) and Context B (black; n=10), revealing qualitatively distinct temporal distributions. (E) Empirical cumulative distribution function of predicted peak times during recall in Context A, with fitted theoretical distributions. Model comparisons favored a Power-law distribution, consistent with temporally compressed structure. Cxt A (n=11; cells=1371) (F) Empirical cumulative distribution function of predicted peak times during recall in Context B, with fitted theoretical distributions. Model comparisons favored a Power-law distribution, closely followed by a uniform distribution, indicating weaker temporal structure than Context A. Cxt B (n=10; cells=924). (G) Variability of astrocytic peak timing across fear conditioning (grey; n = 2746 cells), Context A (blue; n = 1354 cells), and Context B (black; n = 823 cells). Lower values indicate more consistent temporal organization. Points indicate individual astrocytes. Variability increased during recall in both contexts compared to fear conditioning (linear mixed effects model with Tukey HSD). Significance levels: *p ≤ 0.05, **p ≤ 0.01, ***p ≤ 0.001, ****p ≤ 0.0001. See Supplemental Table 1 for statistical details.

At the level of individual animals, eight of the eleven animals exhibited statistically significant Spearman’s rho values relative to their own shuffle-based null distributions and showed significant relationships between true and predicted peak times using the same Huber loss model applied in CFC **(Supplemental Table 1, Extended Data Fig. 14)**. In contrast, of the seven animals that could be analyzed in Cxt B (three animals had <2 detected events), only two showed statistically significant rho values relative to their shuffle-null distributions and significant relationships between true and predicted peak times **(Supplemental Table 1, Extended Data Fig. 15)**. Together, these results indicate that the sequential dynamics observed in Cxt A are more structured than those in Cxt B.

### Parameter analysis further supports sequence organization across spontaneous events

We then applied the same hierarchical Bayesian model used above for the data acquired in Cxt A and Cxt B **(Extended Data Fig. 16)**. We first visualized the estimated peak locations in each context using histograms with smoothed kernel density overlays **(Fig. 4d)**. The two contexts showed clear differences in the distribution of peak locations. In both contexts, we observed bimodal structure, consistent with substantial population-level variability in recalled peak times (Cxt A: Peak distribution median = 6.3s; interquartile= 12.7s; 90^th^ percentile = 23.0s; Cxt B: Peak distribution median = 14.6s, IQR=16.1s, 90^th^ percentile=27.2s). Notably, while both contexts exhibited this bimodality, the second mode was much more pronounced in Cxt B (**Fig. 4d**).

To quantify these distributions, we fit the same set of candidate probability models used in the FC analysis: Exponential, Gaussian, Power-law, and Uniform, and compared predictive performance using WAIC **(Fig. 4e-f)**. In Cxt A, the Power-law provided the best account of the peak-time distribution (lowest WAIC; WAIC = 6914.4) (**Fig. 4e**). In Cxt B, although the Power-law achieved the lowest WAIC (WAIC = 5141.9) **(Fig. 4f)**, its advantage over the Uniform distribution was modest (WAIC = 5184.9), consistent with weaker or heterogeneous temporal organization. Together, the near-tie between Power-law and Uniform in Cxt B, combined with the pronounced bimodality in the peak distribution, suggested that additional structure (beyond a single marginal distribution fit) might be important for disambiguating these models.

We therefore repeated the model comparison after stratifying the data by biological sex **(Extended Data Fig. 17)**. First, we confirmed that the number of detected events was comparable across sexes in both contexts, with no main effects of sex or context and no interaction (two-way ANOVA; sex: F(1,19)=4.84, p=0.100; context: F(1,19)=0.01, p=0.930; interaction: F(1,19)=0.08, p=0.830; **Extended Data Fig. 17a**). We then replotted the peak-location distributions by sex. In Cxt A, males and females showed highly similar peak distributions **(Extended Data Fig. 17b; Supplemental Table 1)**. In contrast, in Cxt B the distributions diverged, with clear differences between males and females that were especially evident in the expression of the second mode **(Extended Data Fig. 17b; Supplemental Table 1**).

We next fit the same candidate distributions to the sex-stratified peak distributions and again found that the Power-law was the best-fitting model by WAIC for both sexes **(Extended Data Fig. 17c)**. However, closer inspection revealed that the apparent Power-law advantage in Cxt B *males* largely arose because, at certain parameter values, the Power-law distribution is practically indistinguishable from the Uniform distribution, rather than reflecting strong evidence for the compressed temporal structure observed in Cxt A. Specifically, the posterior estimate of the Power-law exponent in males was near zero (α=0.19, [0.07 - 0.34]), which makes it difficult to distinguish from a Uniform probability model. Consistently, the Uniform model was essentially tied with the Power-law in Cxt B males (WAIC_PL = 3478.9 vs WAIC_Unif = 3485.5; ΔWAIC ≈ 6.6). In females, α was shifted away from this near-Uniform parameter value (α=0) (**Extended Data Fig. 17d**), making it clear that a Uniform model is less likely given the data, though the Uniform model remained a relatively strong competitor. This interpretation is further supported by the posterior distributions of α across contexts and sexes: in Cxt A, the male and female α posteriors overlap strongly, whereas in Cxt B they are largely non-overlapping **(Extended Data Fig. 17d)**. Thus, whereas Cxt A shows a robust Power-law–like organization consistent with temporally compressed sequences, Cxt B is better characterized as heterogeneous across subjects and sexes, approximately uniform (or weakly structured) in males, with a less uniform and less easily summarized pattern in females, pointing to a different temporal organization during recall in the novel context.

Lastly, we investigated how variable individual peak estimates were across all contexts. Our model gave separate estimates for each individual cells’ peak location, whether driven by shock or occurring spontaneously. This approach allowed us to assess the consistency of temporal information for each cell by estimating an individual standard deviation (SD) parameter for peak locations across events. A lower SD indicates that a cell’s peak activity occurs at a more consistent point within the sequence of events, reflecting greater stability. Based on our non-parametric analyses (**Fig. 4a-b**), we hypothesized that cells from animals re-exposed to the familiar context (Cxt A) would show less variability in their peak locations compared to those in the novel context (Cxt B). Consistent with this prediction, a linear mixed effects model revealed a significant effect of context on peak variability across sequences (**Fig. 4g**). Relative to FC, both Cxt A (β = 2.99, z = 15.58, p<0.0001) and Cxt B (β = 5.56, z = 23.39, p<0.0001) showed increased variability. Importantly, *post hoc* comparisons confirmed that peak variability was significantly lower in Cxt A than in Cxt B (Bonferroni-corrected, p<0.0001). This indicates that astrocytic sequences are more stable in a familiar, conditioned context compared to a novel one, reinforcing the results of our earlier analyses.

Given the evidence for sex stratification in earlier analyses, we repeated the SD comparison separately for males and females. Peak variability did not differ between males and females in Cxt A. In contrast, males exhibited significantly greater peak variability than females in Cxt B, reflected by a significant sex x context interaction (linear mixed effects model; Male x Cxt B: β = 4.27, p<0.0001; **Extended Data Fig. 17e)**. Further, peak variability was reduced during FC relative to Cxt A (β = −3.66, p<0.0001), with a significant male x FC interaction (β = 1.49, p<0.0001). Motivated by these differences, we stratified our cross-validated heatmap analysis by sex **(Extended Data Fig. 17f)**. Both males and females showed significantly higher correlations in Cxt A relative to their shuffled null distributions, whereas neither group showed correlations distinguishable from null in Cxt B (paired and independent *t*-tests), reinforcing the conclusion that temporal organization is reliable in the familiar Cxt A but degraded in the novel context.

### Sex-specific relationship between detected sequences and animal freezing

Our next analysis examined how these detected events from our previous analysis **(Fig. 3d)** are related to animal behavior. Prior work from our lab and others has shown that astrocytic activity is often temporally correlated with behavioral epochs^16,17,28–31^. First, we found that mice did not exhibit substantial freezing in Cxt B (**Fig. 1d**), where robust sequences are also absent, as reported above. In contrast, we observed both robust sequences and substantial freezing for mice who were re-exposed to Cxt A, suggesting a correlation between these robust sequences and fear memories. Then we plotted epochs of freezing on top of the summed activity traces to note the presence (or lack thereof) of freezing in relation to detected sequence events (**Fig. 5a; Extended Data Fig. 12-13)**. Interestingly, in line with our prior observations, freezing was less apparent during sequences. To test this hypothesis, we calculated the average percent freezing during a high activity event and outside of this time-period for animals who were re-exposed to Cxt A and B (**Fig. 5b**). We quantified this relationship by comparing freezing during detected sequence windows to freezing outside those windows, expressed as a within-session difference score Δfreezing = freezing_during_sequence − freezing_outside; negative values indicate *less freezing during sequences.* With a linear mixed effects model, we compared these differences between the two contexts and sexes, finding no differences between sexes (p=0.065), but that there was a sex- and context-specific effect, namely that male mice had less difference in their freezing behavior in Cxt B, regardless of when detected events occurred ([Cxt B x Male]: p=0.010). We could not observe the same interaction for female mice ([Cxt B x Female]: p=0.392) (**Fig. 5b**). This may be due to differences in generalization levels across males and females,^32^ with female mice displaying increased freezing behavior in a novel context, and thus, potentially increasing the presence of sequences in Cxt B. This behaviorally-relevant feature of astrocytic sequences suggests that these events may be memory-specific in nature and preferentially expressed during recall in the conditioned context^10,12,16,17,33^.

**Fig. 5:**
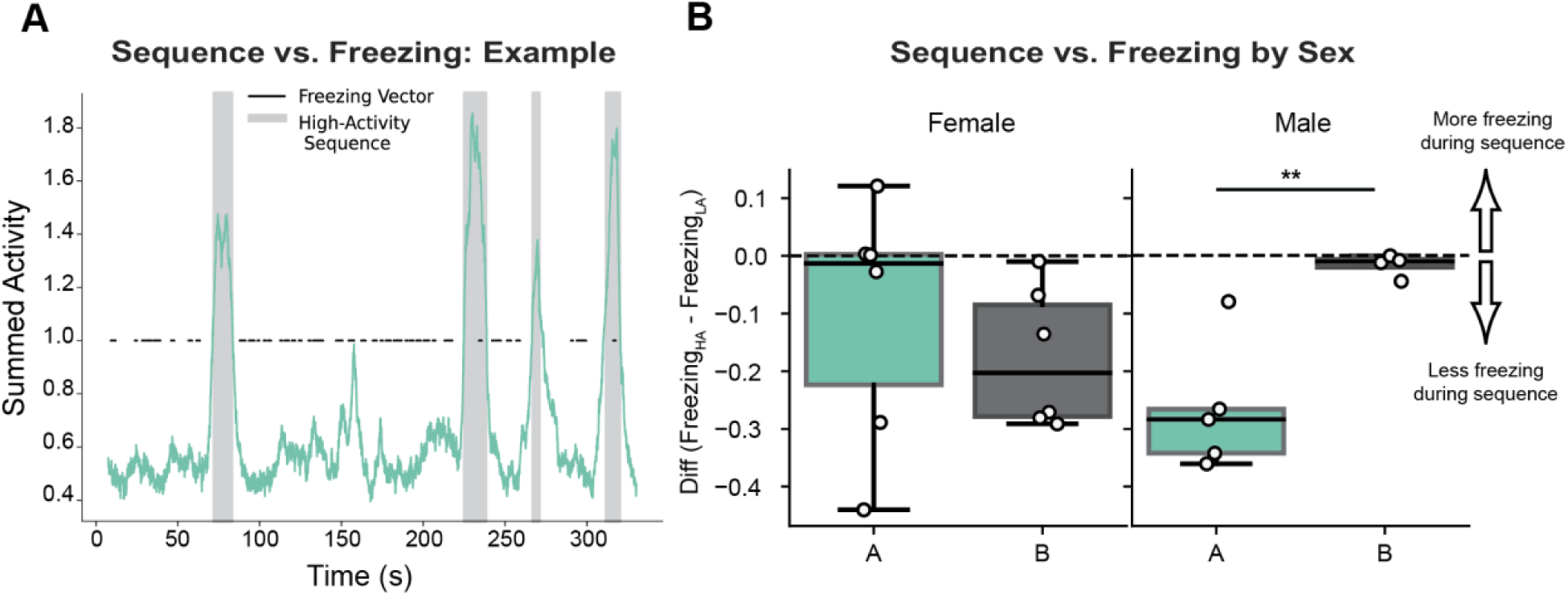
Detected astrocyte sequences display evidence of behavioral relevance in male mice. **(A)** Example summed population astrocytic activity trace during recall in Context A illustrating an inverse relationship between detected epochs of high activity (grey; >1.0 SD peak amplitude and >2.0 second duration threshold) and freezing periods (black dashes). **(B)** Difference in freezing behavior during high activity (HA) ‘sequence’ periods compared to low activity (LA) ‘non-sequence’ periods during recall, calculated as (FreezingHA) - (FreezingLA) by sex and context. Points represent individual animals; boxplots indicate the median and interquartile range, and the dashed horizontal line denotes no difference. A linear mixed-effects model revealed a significant Context × Sex interaction, with male mice showing greater freezing during non-sequence periods in the fear context, an effect not observed in females that is consistent with greater generalization of fear to the neutral context^32,48,49^. Significance level: *p ≤ 0.05, **p ≤ 0.01, ***p ≤ 0.001, ****p ≤ 0.0001. See Supplemental Table 1 for full statistical details.

### Cross-day astrocyte sequence order showed variable preservation across animals

Given the structural similarity between the sequences expressed in Cxt A and those observed during FC, we performed additional analyses to test whether these events relate to contextual memory. We pursued two complementary approaches. First, at the cellular level, we asked whether the temporal ordering evident during FC was preserved in the reactivated population during recall the following day. To assess whether reactivated cells provide a stable substrate for memory, we asked whether animals re-exposed to Cxt A retained a consistent cross-day ordering of activity. Because the proportion of astrocytes active across days did not differ significantly across session pairs (FC:Cxt A or FC:Cxt B) (**Fig. 1h**), we focused instead on differences in *how* these reactivated populations were organized during recall (Cxt A) versus novel context (Cxt B) exposure. We repeated our prior cross-validation approach **(Fig. 2d; 4a-b)**, restricting the analysis to cells active across days and generating cross-validated heatmaps **(Extended Data Fig. 18a)**. Unlike our earlier within-session ordering (Odd sorted by Even), cells were sorted by their peak times measured during FC, and this FC-derived ordering was then applied to activity during the context session (Cxt A or B) to assess cross-session stability. Qualitatively, the progression appeared more structured in Cxt A than Cxt B **(Extended Data Fig. 18a, Extended Data Fig. 19)**.

Quantitatively, however, evidence at group level was mixed when we compared observed Spearman’s ρ values against shuffle-based null distributions, neither context showed a significant elevation above shuffle (paired t-tests: [Cxt A] t=1.60, p=0.140; [Cxt B] t=-0.17, p=0.868; **Extended Data Fig. 18b**). Consistent with this, null-corrected correlations (observed minus null mean) did not differ between Cxt A and Cxt B (independent t-test: t=1.32, p=0.204; **Extended Data Fig. 18b**). Because the Cxt A heatmaps nonetheless suggested meaningful ordering in several animals, we next tested significance at the individual level using a bootstrap procedure as in our prior analyses (**Supplemental Table 1**). At the animal level, three of six Cxt A animals showed significant cross-day sequence correlations (after multiple-test corrections; Benjamini-Hochberg; FDR=0.01), whereas none of the six Cxt B animals did **(Extended Data Fig. 18c)**, indicating heterogeneity that is not well captured by the pooled group comparisons. To further evaluate these individual-animal results, we tested whether the observed number of significant animals in each context exceeded that expected under a binomial null model with success probability *p* = 0.05. The proportion of significant animals in Cxt A was greater than expected by chance (exact binomial test, *p* = 0.002; **Extended Data Fig. 18c**), whereas the proportion in Cxt B was not (*p* = 1.0). Nevertheless, the proportions did not differ significantly between contexts when compared directly using Fisher’s exact test (*p* = 0.182; **Extended Data Fig. 18c**).

Finally, motivated by our prior results, we stratified the analysis by sex. This analysis was limited by attrition (animals lacking sufficient events in one context) and by exclusion of animals with fewer than 10 reactivated cells; as a result, only two females contributed data. As expected given the low power, we detected no differences in females (paired t-tests: [Cxt A–F] *t* = 0.67, *pBH* = 0.625; [Cxt B–F] *t* = 1.46, *p* = 0.382; Extended Data Fig. 18d). In males, neither context differed individually from shuffle ([Cxt A–M] *t* = 2.79, *pBH* = 0.102; [Cxt B–M] *t* = −0.66, *pBH* = 0.559). A direct comparison of null-corrected correlations between contexts in males yielded an uncorrected *p* = 0.049, but this effect was not significant after Benjamini–Hochberg correction (*pBH* = 0.102*; Extended Data Fig. 18d). These analyses therefore suggest that any sex-specific cross-day ordering effects should be interpreted cautiously.

### Contextual fear conditioning is necessary to explain later changes in the fear context

A key question was whether the effects observed in Cxt A depended on prior fear conditioning, or whether similar patterns would be present in the absence of shock (i.e. during baseline exposure to the same environment). Our behavioral paradigm includes an initial habituation session (**Fig. 1a**), which occurs in the same physical context that later becomes associated with fearful stimuli, providing an ideal control condition. We therefore analyzed habituation recordings using the same workflow as in our primary analyses.

We first plotted representative activity traces from all animals during habituation **(Extended Data Fig. 20a)**. Visual inspection did not reveal clear or consistent patterns of population activity, so we next tested whether the number of detected events differed between habituation and subsequent sessions using the event detection method described in **Fig. 3d**. We observed a significant difference in the number of events between habituation and Cxt A recall (Kruskal–Wallis, *p* = 0.018; Mann–Whitney [Cxt A–HAB], *p* = 0.022). In contrast, we found no difference between habituation and the subsequent exposure to Cxt B (*p* = 0.138), nor between Cxt A recall and Cxt B (*p* = 0.266) **(Extended Data Fig. 20b**). This suggests that astrocytic population activity increases in Cxt A specifically following conditioning.

To assess any temporal organization of events during habituation, we computed Spearman correlations between successive events as in our prior analysis. However, only two animals exhibited more than a single event during habituation, limiting group-level analysis. We therefore compared each animal individually against its own shuffle-based null distribution. One animal showed significant event-to-event correlation **(Extended Data Fig. 20c),** whereas the other did not **(Extended Data Fig. 20d).** This variability, combined with the low event number, indicates an absence of robust or consistent sequential organization during habituation. Finally, we fit the same hierarchical Bayesian model used previously to the detected event astrocytic calcium dynamics for these two animals. As in earlier analyses, the Power-law model provided the best fit. However, consistent with what we observed in Cxt A males **(Extended Data Fig. 17d)**, the estimated Power-law exponent was close to uniform-like behavior (α = 0.28, 95% CI = [0.07, 0.48]; WAIC_Power Law = 1189.1; WAIC_Uniform = 1192.9; **Extended Data Fig. 20e**), suggesting weak temporal structure during habituation. Together, these results indicate a clear lack of robust sequential activity in the conditioning context prior to unconditioned stimulus (US) presentation, supporting the interpretation that the temporal organization observed in Cxt A recall emerges after learning rather than reflecting baseline properties of Cxt A exposure.

## Discussion

Our findings identify astrocytes in dorsal CA1 as a source of robust, temporally organized calcium dynamics during both the acquisition and retrieval of contextual fear memory. These dynamics are strongly engaged by salient experience and exhibit statistical features that parallel key properties of hippocampal time-cell representations. During conditioning, astrocytic population activity is reliably engaged by foot shocks, expressing temporally compressed post-shock sequences that are stable across stimulus presentations. During recall, spontaneous high-activity events are observed in both contexts; however, only animals returning to the conditioned context (Cxt A) show consistent evidence of sequential organization. In both contexts, a subset of astrocytes remains active across days. Together, these results demonstrate that astrocytic population activity can exhibit temporally structured dynamics linked to aversive learning and memory recall.

The analytical framework developed here may add interpretive power by revealing latent temporal structure that is obscured by population-averaged measures. For example, previous work has shown that hippocampal astrocytes exhibit anticipatory ramping in calcium activity preceding an expected reward^8^. An important possibility raised by our results is that such population-averaged ramps may reflect an underlying sequential organization that becomes apparent when activity is analyzed at the level of cross-validated ordering and peak-time distributions, rather than through mean activity alone. Similarly, other studies have demonstrated that spatial position along a linear track can be decoded from hippocampal astrocytic calcium activity dynamics in a manner analogous to neuronal place fields^7^. Notably, these data reveal sequence-like activity across the track that ramps in response to reward delivery, a phenomenon also observed in canonical hippocampal time cells and consistent with the above mentioned study^6–8^. Together, these observations suggest that temporally structured astrocytic population dynamics may represent a general feature of hippocampal processing in tasks with temporal contingencies, warranting future investigation into their role in memory-guided behavior.

We posit that astrocyte population sequences encode temporal context information in a manner that parallels hippocampal time cells. This interpretation is supported by work examining how hippocampal neurons process time during tone conditioning paradigms^33–35^. Across studies, neuronal activity is observed around the tone–shock association, with subsets of neurons exhibiting time-locked responses at specific moments within the trace interval, sequential activation patterns emerging across trials, and neurons with shared temporal tuning showing elevated spontaneous correlations even prior to learning. Consistent with this framework, several statistical features of the astrocytic sequences we identify, including temporal-compression, align with this interpretation and represent an important direction for future investigation^21^.

At the same time, key differences distinguish astrocytic dynamics from canonical neuronal time-cells. Neuronal time cell activity can persist over longer timescales (e.g., minutes), whereas astrocytic sequences are shorter-lived. This suggests that astrocytes may contribute complementary temporal information and integration processes rather than directly mirroring neuronal coding^6,36^. This idea is in line with recent work that did not observe clear sequential activation in the hippocampus in a similar paradigm, yet they could decode memory-relevant stimulus identity from longer-timescale neuronal population dynamics^34^. Mnemonic information can thus be encoded even without canonical sequences, raising the possibility that astrocytic dynamics participate in a parallel temporal coding schema, operating on distinct timescales. Several methodological factors may also contribute to these differences in characteristics. One-photon calcium imaging provides lower spatial resolution than two-photon approaches, potentially limiting our ability to fully resolve astrocytic subdomains and weaker late events. Consistent with this, later calcium events exhibit reduced signal-to-noise ratios, suggesting that some activity may fall below detection thresholds. In addition, commonly used pipelines for calcium imaging analysis are optimized for neurons and perform sub-optimally for astrocytes, particularly for somatic-specific event detection^37^. Although other astrocyte-specific tools exist, they often emphasize process-level activity or require manual somatic identification^38^. Given this, we manually segmented astrocytes based on morphology and prominent somatic events to enable cross-day tracking, which signals the need for tools developed for astrocyte soma specific identification. Even so, these technical constraints may have reduced sensitivity to weaker or spatially-distributed events, contributing to the variable evidence for cross-day coupling observed in our work. Higher-resolution, cell-type-optimized imaging and analysis approaches will be necessary to fully resolve the stability and structure of putative astrocytic temporal dynamics across days. An additional interpretive consideration is that stronger sequential organization observed in the conditioned context during recall may not reflect memory-specific temporal coding alone. Because the conditioned context is also the acquisition environment, differences between Cxt A and Cxt B could arise from contextual similarity, familiarity, or generalization-related processes. Although the no-shock behavioral variant partially dissociates exposure order and context experience in the absence of reinforcement, it does not fully exclude the possibility that sequence-like astrocytic population dynamics may also differ across contexts independent of associative learning.

Recent work positions astrocytes as integral components of memory-related ensembles rather than passive correlates of neuronal activity. Experience can recruit astrocyte ensembles whose reactivation is sufficient to promote recall and amplify neuronal engram reactivation^13^. Complementary studies suggest that astrocytes can form multi-day eligibility traces that are preferentially engaged during recall via enhanced neuromodulatory responsiveness^14^. In parallel, calcium imaging has revealed centripetal integration from astrocytic processes to soma gated by norepinephrine, providing a plausible mechanism for astrocyte state transitions^39^. These findings provide a framework for interpreting our findings: salience-linked neuromodulation may recruit temporally compressed astrocytic sequences during conditioning and recall^40–44^. The behavioral association we observe, reduced freezing during detected sequence windows (particularly in males), further supports the idea that these events coincide with state transitions relevant for fear expression. This model generates several testable predictions. First, simultaneous astrocyte-neuron could clarify temporal coupling; if astrocyte sequences scaffold neuronal dynamics, then trial-to-trial variability in astrocyte sequence stability should predict variability in neuronal sequence structure and recall-related ensemble activity. It will be informative to extend this approach to more complex trace conditioning paradigms to determine if these astrocytic sequences are also present during tone-delay-shock pairings, and how they compare directly with established neuronal time cell dynamics. Second, causal perturbations of astrocytes during conditioning or recall should alter the subsequent organization of recall events, potentially degrading context-specific sequential structure and disrupting fear expression. Such manipulations would help determine whether astrocytic sequences are necessary components of memory-state dynamics or instead reflect downstream activity from neuronal circuits. Third, higher-resolution imaging, including dual-cell-type or two-photon approaches, will be essential to resolve subcellular dynamics and process-to-soma integration. Finally, if norepinephrine gates astrocytic sequences, then reducing adrenergic signaling should selectively weaken sequence structure during conditioning and recall^6–8,13,14,21,35–47^.

In summary, astrocytic population activity in dorsal CA1 exhibits robust, temporally organized dynamics that are engaged by salient experience and expressed during memory recall. These astrocytic sequences share key statistical features with hippocampal time-cell representations but operate on distinct timescales and show heterogeneous cellular stability across days. By revealing sequence-level organization that is not captured by population-averaged activity, this work provides a unifying perspective to reconcile prior observations of astrocytic activity. However, given the correlational nature of the data and the possibility that detected events reflect broader behavioral state transitions, the specificity of these dynamics to memory remains unanswered. Future studies combining neuronal recordings, causal astrocyte perturbations, and higher-resolution imaging will be required to determine how astrocytic and neuronal dynamics interact to support memory-guided behavior.

## Supporting information

Stats Table

## Author Contributions

R.A.S., R.L.S., and S.R. conceptualized the experimental paradigm. R.A.S., R.L.S., S.C. and E.R. performed animal surgical procedures and calcium imaging experiments. R.A.S., R.L.S., R.C., E.R., and A.M. performed data analysis. M.D.B. and R.L.S. performed behavioral analysis. R.A.S, R.L.S., and R.C. generated figures. R.C. performed Bayesian Modeling. R.A.S, R.L.S., R.C., and S.R. wrote the paper. R.A.S., R.L.S., R.C., M.H., and S.R. interpreted the results and analysis. R.C. and M.H. provided expertise on temporal-sequence analysis.

## Acknowledgements

This work was supported by the NIH Transformative Award (NIH R01DE033519), the Ludwig Family Foundation, the Air Force Office of Scientific Research award (FA9550-21-1-0310), the Pew Scholars Program in Biomedical Sciences, the Chan Zuckerberg Initiative, the Neurophotonics Center at Boston University and the Center for Systems Neuroscience. Behavioral schematics were created using BioRender. We thank members of the Ramirez Lab, especially Heloise Leblanc, for her helpful feedback on the manuscript.

## Data Availability Statement

All raw data (e.g., original calcium imaging videos) and pre-processed data used to generate figures (e.g., CSV files) will be available upon request to the corresponding author.

## Code Availability

Code used in the current study is available at https://github.com/rsenne/dCA1_Paper.

## Methods

### Subjects

The animals used in this study were male and female, wild-type C57BL6 (P29-35; 17-19g) mice obtained from Charles River Laboratories. Animals were housed in the Boston University animal facilities in standard ventilation cages on a 12:12 dark/light cycle (0700-1900) and given ad libitum access to standard food and water. Mice were housed with littermates in cages with 4-5 mice and after GRIN lens implantation, they were individually housed to prevent any damage to their implants. Following GRIN lens surgery, mice were given 10 days to recover before the start of behavior. All subjects were treated in accordance with protocol 201800579 approved by the Institutional Animal Care and Use Committee (IACUC) at Boston University. No statistical methods were used to determine sample size but were instead based on sample sizes of previous 1p imaging studies.

### Stereotaxic Surgery

#### Viral transduction

Mice were anesthetized with 3-3.5% isoflurane during induction and maintained at 2-3% inhalation using stereotaxic nose cone delivery (oxygen 1.2 L/min). Paralube ophthalmic ointment was applied regularly to the eyes to maintain adequate lubrication. Scalp hair was removed using Veet or Nair hair removal cream; this was subsequently removed using applications of betadine solution and ethanol (70%). Lidocaine hydrochloride (2.0%) was injected subcutaneously (SQ) as a local analgesic agent under the scalp prior to incision. Meloxicam (0.1mg/kg) was administered SQ at the start of surgery. After scalp incision, mice received a unilateral (right) craniotomy with a 0.5-0.6 mm drill bit for the dorsal hippocampal CA1 (dCA1) viral injection. A 10uL airtight Hamilton syringe with an attached 33-gauge beveled needle was used to inject AAV-GfaABC1D-cyto-GCaMP6f-SV40 (Penn Vector Core) at the coordinates of dCA1: -2.00 anteroposterior (AP), +1.40 mediolateral (ML) and -1.50 dorsoventral (DV). All coordinates are given relative to bregma (mm). 1000nL of undiluted virus was injected at 50nL/min using a micro infusion pump (UMP3; World Precision Instruments). After injection, the needle remained at the site of injection for 5-7 minutes to prevent viral spread. After injection, the incision was sutured closed using 4/0 Non-absorbable Nylon Monofilament Suture (Brosan). At the end of surgery, mice were injected with a 0.1mg/kg intraperitoneal (IP) dose of buprenorphine. Following surgery, mice were placed in a clean recovery cage with a heating pad until they regained reflexes. To allow time for viral expression and recovery, mice were monitored for 2 weeks prior to implantation of GRIN lens surgery.

#### GRIN lens implantation

After 2 weeks, we inserted a GRIN lens (Inscopix, Inc.). 30 minutes prior to surgery, mice were injected with 0.1mL of dexamethasone diluted in 0.1mL sterile saline IP to reduce brain swelling. Mice were prepared as described above, and the previous incision site was reopened. A 0.5-0.6 mm drill bit was used to drill a 1 mm craniotomy around the previous site of injection at the coordinates listed above. Using vacuum suction, 30 to 34-gauge needle attachments and constant administration of cold, sterile saline, brain tissue was aspirated down just beyond the thick fibers of the corpus callosum. Blood control was performed using an absorbable gelatin sponge (Surgifoam). A GRIN lens and integrated base plate were lowered slowly to -1.30 DV and a biocompatible silicone adhesive (Kwik-Sil) was used to seal off the exposed perimeter of the lens and skull. Multiple layers of adhesive cement (C&M Metabond) were applied up to the cuff of the integrated base plate and the headcap was completed with darkened dental cement (Stoelting). Mice were administered the same doses of postoperative analgesics described above. They recovered for 10 days before imaging experiments were performed during our behavioral paradigm.

### Behavioral testing

Prior to behavioral testing, all mice were handled and habituated to being plugged into the miniature microscope. On Day 1, mice were subjected to contextual fear conditioning (FC) for 330s in Context A (Cxt A), where they received foot shocks (0.75mA, 2s duration) at the 120, 180, 240 and 300s time points. On Day 2, mice underwent a 330s contextual recall session in the same mouse conditioning chamber (Cxt A) or were instead exposed to a novel Context B (Cxt B) for the same duration of time (330s).

### Context parameters

Cxt A was a standard mouse conditioning chamber (18.5 x 18.5 x 21.5 cm; Coulbourn Instruments) with metal-panel side walls, plexiglass front and rear walls and a stainless-steel grid-floor (16 bars) that were connected to a precision animal shocker (Actimetrics). Cxt B was a standard mouse conditioning chamber of the same size in a different behavioral room that was modified to have striped plastic walls, almond scent, and a transparent plastic floor. No lighting differences or the presence of auditory stimuli existed in either context. Ethanol (70%) was used to clean Cxt A, while Rescue disinfectant spray was used to clean in Cxt B to maintain these odor profiles.

### One-photon Imaging

#### Acquisition

All imaging experiments were conducted on awake, behaving mice in a freely moving manner in standard mouse conditioning chambers. GCaMP6f fluorescence signals were imaged using a miniature integrated fluorescence microscope system (Inscopix Inc.) with GRIN lenses implanted in the dorsal CA1 of the hippocampus (dCA1). Before the start of imaging, the miniature microscope was attached to the baseplate and adjusted to determine the best focal plane and gain for each mouse to achieve the best fluorescence signals. These values were maintained across all recording sessions. Frequency (Hz) and LED output power were consistent across all mice. Behavioral video was captured using the nVista Imaging System (Inscopix Inc.) that was triggered by an external TTL to time lock our behavioral session to calcium recordings (FreezeFrame4).

#### Pre-processing

Calcium imaging data were pre-processed using the Data Processing Software (Inscopix, Inc.; IDPS). Videos were cropped from 1280 x 800 pixels to 1200 x 800 pixels and spatially down sampled by a factor of 4, from 1200 x 800 pixels to 300 x 200 pixels and temporally down sampled by a factor of 2 from 20 to 10 Hz. A spatial bandpass filter was applied using the default settings in IDPS (low cut off: 0.005 and high cut-off = 0.500). Motion correction was performed with the first frame as the global reference frame origin. PCA-ICA and CNMFe analysis failed to identify individual astrocytes in a reliable manner. Thus, to identify individual cells, pre-processed images were converted to dF/F using the mean frame as a reference and ROI selection was performed within IDPS. For each animal and session, the field of view was visualized as both the mean and maximum projection across all frames to identify active cells. Individual ROIs were drawn around pixel groupings that showed clear fluorescence modulation across time, determined by scrolling through each frame of the dF/F movie and comparing it to the projection images. A region was considered an ROI if it met the following criteria: (1) Spatially distinct: round or oval in shape with clear boundaries from adjacent cells; at our resolution, no processes could be visualized. (2) Visible activity: Displayed visible fluorescence fluctuations or calcium transients in the dF/F trace extracted; low-SNR structures were excluded. (3) Stability: The structure remained stable throughout the session after motion correction. Using these guidelines, ROIs were drawn freehand using the polygon selection tool to match the cell boundaries (e.g., a hexagon shape was drawn within each soma). To ensure consistency across animals, ROI selection was performed and cross-validated across multiple experimenters who were blinded to animal identity and session. Any ambiguous structures were excluded at this step. Following ROI selection, all single-astrocyte traces were further evaluated for signal quality while visually inspecting each ROI.

#### Longitudinal registration

Longitudinal cell ROI registration was performed across imaging sessions using the open-source Matlab toolbox program, CellReg, which utilizes a probabilistic framework method to identify the same cells for tracking cells across multiple recording sessions^15^. In brief, it first aligns each session field-of-view (FOV) to one another by projecting the centroid locations of all the cells’ spatial footprints onto a single one image and maximized the cross-correlation between each session’s projection and a reference session (Day 1; fear conditioning). We utilized non-rigid transformations, as they best aligned our dataset. Following spatial alignment, CellReg computed Pearson correlation coefficients between spatial footprints of neighboring cells across sessions. These correlations, together with the centroid distances (max distance threshold: 13-14 um), were used to estimate the probability that two ROIs were the same cell across days (Psame). The resulting data were modeled as a weighted sum of the two distributions corresponding to same-cell and different-cell pairs, allowing for estimation of false-positive and false-negative registration rates. A Psame threshold of 0.50 (default) was used as input to a clustering algorithm to generate a final cell registration assignment.

Throughout the registration process, intermediate outputs were visually inspected to ensure optimal parameter selection and alignment quality. This included examination of raw spatial footprint overlaps, spatial footprints following linear or non-linear transformations, outputs of probabilistic modeling based on centroid distance or spatial correlation thresholds and overlaps of final registered spatial maps (see the original CellReg paper for visualizations of these steps). Further, registered cell outputs were exported and further validated using custom Python scripts, which enabled manual inspection of every registered cell pair for each session comparison (i.e., HAB:FC, FC:A, FC:B). These scripts also generated CSV files containing the indices of each cell on Day 1 and Day 2, with a value of –1 indicating that a given cell was not detected in a particular session. If registration quality was determined to be suboptimal at any step, either (1) an affine-transformed version of the raw spatial footprint was used to improve initial alignment prior to CellReg or (2) the cell pairing for that session was manually excluded prior to analysis.

While this approach is constrained by the capabilities of CellReg and relies on visual assessment by multiple blinded experimenters, best efforts were made to ensure rigorous and transparent registration and ROI detection, given the current lack of automated astrocyte-specific somatic detection tools. Importantly, CellReg has been successfully applied in numerous prior studies across cell types, supporting its suitability for tracking somatic astrocytes across days^50,51^.

#### Behavioral analysis

Freezing behavior was scored using Any-Maze, using a bout threshold of ≥ 1.25 seconds. This allowed for identification of the onset and offset of freezing epochs, as well as quantification of total percent time freezing (%) across the entire session or within defined time-bins (every 60 seconds) (Stoelting Inc.).

### Immunohistochemistry

After the completion of experiments, mice were overdosed with 3% isoflurane and transcardially perfused with cold phosphate buffered saline (PBS), followed by 4% paraformaldehyde (PFA) in PBS. Brains were extracted and placed in PFA at 4℃ for 24-48 hours. All brains were sliced coronally at 50um on a vibratome into cold PBS or 0.01% sodium azide in PBS for long term storage. Native green fluorescent (GFP) protein signal from the GCaMP6f expression was assessed, and sections were counterstained for both astrocytes and neurons using immunohistochemistry (IHC).

For IHC, dorsal CA1 (dCA1) sections underwent three washes in PBS for 5-10 minutes each to remove the 0.01% sodium azide storage solution. Sections were washed in 0.2% Triton X-100 in PBS (PBST) for 3 x 5 minutes each at room temperature. Next, slices were blocked for 2 hours in 1% bovine serum albumin (BSA) and 0.2% Triton X-100 in PBS solution on a shaker at room temperature. Sections were incubated in primary antibodies (1:1000 mouse anti-GFAP [NeuroMab] or 1:500 polyclonal guinea anti-NeuN/Fox3 [SySy]; 1:1000 chicken anti-GFP) made in the same 1% BSA/PBS/Triton X-100 solution at 4C for 24-48 hours.

Sections underwent 3 x 5-minute washes in PBST to remove the primary antibody solution. Sections were incubated in secondary antibodies (1:1000 Alexa Fluor 555 goat anti-mouse IgG [Invitrogen] or 1:1000 Alexa Fluor 555 goat anti-guinea pig [Invitrogen]; 1:1000 Alex Fluor 488 goat anti-chicken [Invitrogen]) for 2 hours at room temperature on a shaker. Sections were washed 3 x 10 minutes in PBST, and subsequently mounted onto microscope slides (VMR International, LLC) with Vectashield HardSet Mounting Medium with DAPI (Vector Laboratories, Inc.). Slides were allowed to dry for 24 hours at room temperature and edges of the coverslips were clear nail polished to maintain moisture. Slides were stored in a slide box in 4C after confocal imaging. Any mice that had visible astrogliosis were removed from subsequent analysis (e.g., visual assessment of GFAP intensity increase, thickening of soma, and processes and proliferation of cells towards the site of damage (window area)).

To quantify cellular specificity of the astrocyte-specific GCaMP6f construct, we performed the above immunohistochemical analysis of the dCA1 pyramidal cell layer underneath the GRIN lens. Images were acquired on a Zeiss LSM 800 epifluorescence microscope using the Zen 2.3 software. Here, we imaged 1 x 3 tile regions unilaterally for each animal at 20x magnification and a z-stack with 2µm step size. Images of DAPI (blue), GCaMP6f (green) and GFAP or NeuN (red) were acquired with these specifications, they were post-hoc split by channel in FIJI (ImageJ) and average projected for the green and red channels. Using Cellpose, an algorithm for cell segmentation, we identified the total number of GCaMP6f+ somas in each image and saved the resulting cell masks into FIJI format^52^. These masks were overlaid on the GFAP or NeuN/GCaMP6f overlap images to identify the number of overlapping (yellow) cell masks. The specificity was quantified as the number of GFAP or NeuN cells co-expressing GCaMP6f (yellow) divided by the total number of GCaMP6f+ cells within that image. Each mouse has a resulting average that is representative of n = 3 slices per animal that is included in the plot (**Extended Data Fig. 1**). Representative images of these overlaps (GFAP/GCaMP6f or NeuN/GCaMP6f) are included in this figure as well.

### WaveMAP Clustering

Exploratory WaveMAP analysis of astrocytic calcium waveforms during contextual fear conditioning (FC) were performed following the protocol described in and Extended Data Fig. 8 and generated using code adapted from the WaveMAP_Figures_Data.ipynb Jupyter notebook (https://github.com/EricKenjiLee/WaveMAP_Paper), as well as custom code generated by Senne & Suthard for visualization.^26,27^ First, we compiled all single-astrocyte calcium traces from all animals (n = 14) during FC and normalized the data via z-scoring. Next, for every trace, we generated an average shock response [120, 180, 240, 300 second time points] with a window of –7.5 seconds to 15 seconds after shock. Here, we concatenated these average shock responses for all animals into a NumPy array that was fed into WaveMAP. This NumPy array consisted of the time (column; 3303 indices) and cell row (2823 single astrocytes) across all individual animals within FC. Broadly, WaveMAP first normalizes the array by dividing each value by the maximum value in each row using NormWF. Next, it passes NormWF into UMAP to generate a high-dimensional UMAP graph. It further applies Louvain clustering to the graph and allows visualization of the detected clusters within UMAP space. The WaveMAP parameters were optimized and evaluated using a classifier’s performance; n_neighbors=30, min distance = 0.1 (**Extended Data Fig 8c**). Once the resulting clusters were optimally confirmed with the classifier, we examined all the individual cluster average waveforms and determined the total number and percentage of astrocytes within each cluster (**Extended Data Fig. 8e**). Once these results were confirmed, we took each of these clusters and plotted the average z-scored calcium waveform peri-event within the –7.5 to 15 second window surrounding foot shock to understand if different functional subpopulations existed in response to fearful stimuli (**Extended Data Fig. 8b,d**).

### Statistical Methods

All statistical analyses were conducted using Python and GraphPad Prism v10. Details of the statistical tests, including the type of test used, sample sizes (n values), test statistics, p-values, post-hoc multiple comparisons, and any data removals, and whether testing met statistical criteria (outliers, normality and variance) are provided in the text, figure legends and the Supplemental Table 1, as applicable.

### Shuffle Based Hypothesis Testing

Many analyses of the current work implement shuffle-based hypothesis testing **(e.g.,** **Fig 2e).** Unless otherwise noted, shuffle-based hypothesis testing was performed using a permutation procedure across cells. A test statistic (Spearman correlation) was first computed from the empirical data, denoted 𝑇_emp_, quantifying the relationship between two vectors of per-cell summary features (e.g., argmax times derived from odd vs. even event splits).

To generate a null distribution, we disrupted the correspondence between cells by randomly permuting one of the vectors (e.g., the “even-sorted” argmax values) across cells while keeping the other fixed. This preserves the marginal distributions of both variables but destroys any cell-specific pairing structure. For each shuffle iteration, the same statistic was recomputed, yielding 𝑇_null_.

Repeating this procedure across many iterations produced an empirical null distribution for the statistic of interest. Statistical significance was assessed by comparing 𝑇_emp_ to this null distribution. Two-tailed p-values were computed as the proportion of shuffled statistics whose absolute value exceeded that of the empirical statistic, with a standard finite-sample correction:

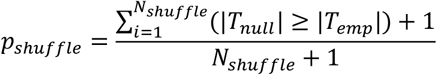

### Normality and Outlier Testing

To ensure the validity of our analyses, data normality for One- and Two-way ANOVAs and Paired- or independent t-tests was evaluated using the Kolmogorov-Smirnov test. Equality of variance was assessed using the Brown-Forsythe test or Spearman’s test for heteroscedasticity. Outliers were identified using the ROUT method, as recommended by GraphPad Prism. This method relies on nonlinear regression to detect outliers, with a ROUT coefficient Q value set at 10% to increase the power of detection while maintaining a relatively lenient threshold. For linear mixed effects models, beta values and group variance are reported within Supplemental Table 1.

### Peri-Event Analysis

For event-triggered average significance (foot shock), we used a tCI confidence interval method previously proposed.^53^ For our event triggered averages, we aggregated all within-animal signals so that we could assume that our samples were independent and identically distributed. Thus, for this method we assumed each time point was distributed according to a student-t’s distribution. We then marked any period greater than 1.0 s (this decision was arbitrary and could be chosen to be longer for more conservative estimation, but this is longer than the proposed time threshold in the original paper) that did not include the baseline of 0 (traces were median-shifted to zero) was marked as a significant peri-event. To quantify shock responsiveness, we compiled all individual astrocyte calcium traces for the entire session from all animals (n = 2823 cells) and grabbed epochs surrounding each foot shock [120, 180, 240 and 300 sec] that included a 5 second baseline and a 15 second post-shock period. We classified a cell as ‘shock-responsive’ if its post-shock peak exceeded 1.96 SD above the baseline (outside of the 95% CI) for at least one shock. This was visually assessed over all cells to ensure accurate detection (see below for a small subset of these; not included in the manuscript). Based on these criteria, 2766/2823 (98.0%) were shock-responsive, while 57/2823 (2.0%) were not.

### Hierarchical Bayesian Model

The hierarchical Bayesian model we implement here assumes each cell increases its activity from baseline level 𝑎_0_according to its Gaussian-shaped temporal receptive field. To account for the variability across sequences, we assume that the peak location 𝜇𝑖 is estimated separately for the 𝑖^𝑡ℎ^ sequence while the field width of 𝜎 and height of 𝑎_1_ are fixed across sequences. Therefore, at each time t after the onset of a foot shock (fear conditioning) or high internal activity (detected by event detector), the expected activity 𝜆 is:

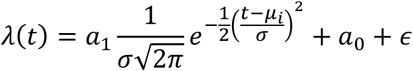

where the small Gaussian error term 𝜖 is centered at zero with a small fixed standard deviation calculated as three times the standard deviation of the entire session for each cell. In addition, we assume that 𝜇𝑖 is sampled from a normal distribution centered at 𝑀 that captures the overall central tendency of the peak and standard deviation 𝜎𝑡 that estimate the between sequence variability to allow for a more accurate estimation of the temporal receptive field width^21^.

The posterior distributions of estimated parameters were generated through the rStan package with four independent Markov-Chain-Monte-Carlo (MCMC) chains (3750 warm-up iterations and 250 post warm-up samples in each MCMC chain). The mode of the posterior distribution was then used to perform additional statistical analysis in the Result section.

### Movement Analysis

To compute instantaneous velocity, we extracted two-dimensional (x, y) position coordinates from DeepLabCut–scored top-down videos^54^. This method was confirmed for accuracy by a blinded researcher that manually scored a subset of videos. Frames with low pose-estimation confidence were excluded using a likelihood threshold, and short gaps were interpolated. The resulting position traces were smoothed using a Savitzky–Golay filter, which enabled computation of analytical derivatives of the locally fitted polynomial. Velocity was then estimated on a frame-by-frame basis as the Euclidean norm of the temporal derivatives of the x and y position coordinates.

To assess the relationship between velocity and astrocytic population activity, we applied principal component analysis (PCA) to z-scored astrocytic activity, yielding a low-dimensional representation of population dynamics. PCA was used to capture dominant patterns of shared activity across astrocytes while reducing noise and redundancy in the full population.

For each principal component (PC), we computed the Pearson correlation coefficient between the PC time series and the animal’s velocity trace. To quantify how strongly velocity was embedded within the population activity space, we defined a *subspace score* as:

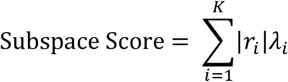

In short, we sum the products of the i-th absolute PC correlation with velocity (|𝑟_𝑖_|) and the i-th PCs explained variance (𝜆_𝑖_). In essence, this can be interpreted as how much population variance lies in the direction of activity space that tracks velocity. To assess statistical significance of the subspace score, we generated a null distribution by circularly shifting the velocity trace relative to neural activity, preserving the autocorrelation structure of both signals while destroying their temporal alignment. The subspace score computed from the real data was compared against this null distribution.

In addition to subspace alignment, we evaluated how well astrocytic population activity could *predict* velocity using decoding analyses. Specifically, we trained a ridge regression model to decode velocity from the top ten PC scores. Ridge regression was chosen to mitigate overfitting and account for multicollinearity among predictors. Model performance was assessed using k-fold cross-validation, and the regularization parameter was selected automatically within the cross-validation framework as implemented in scikit-learn^55^. Decoding performance was quantified using cross-validated coefficient of determination (R²). Statistical significance was again assessed using circularly time-shifted velocity traces to generate a null distribution of decoding performance. Together, these analyses allowed us to characterize both the geometric embedding of velocity within astrocytic population activity and the extent to which population dynamics could be used to predict ongoing movement.

Lastly, we computed the single cell correlations between each individual cell and the respective velocity traces as well as the Pearson correlation between the top PC and velocity.

---

**Extended Data Fig. 1.**
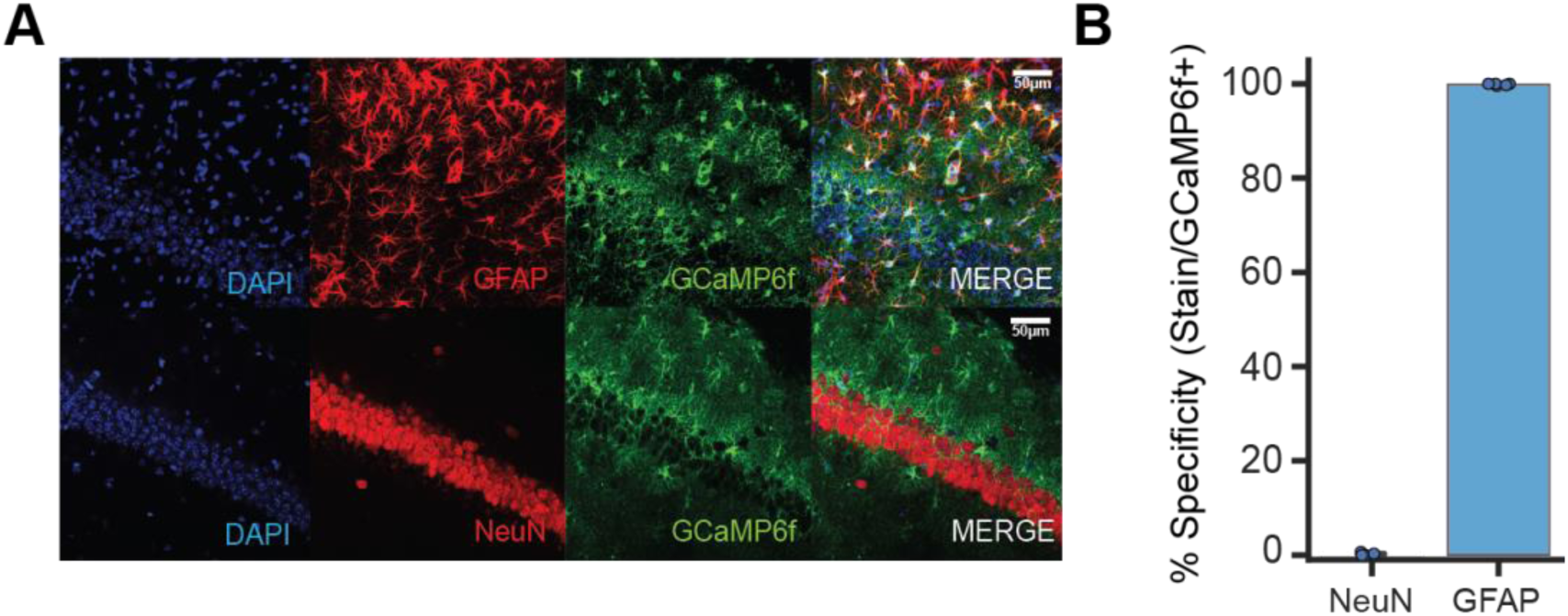
: Specificity of astrocyte viral construct. **(A)** Representative confocal microscopy images of the dCA1 pyramidal cell layer visualizing DAPI+ cells (blue; nuclei), GfaABC1D-GCaMP6f (green; viral expression), glial fibrillary acidic protein (GFAP) (red; astrocytes) or NeuN (red; neurons) to ensure selective expression of our virus in astrocytes at 20x magnification. Scale bar, 50µm. **(B)** Specificity of GCaMP6f (3 NeuN/1326 GCaMP6f = 0.23%; 1204 GFAP/1206 GCaMP6f = 99.83%) with n = 4-5 mice and n = 3 slices per mouse. See Supplemental Table 1 for statistical details.

**Extended Data Fig. 2.**
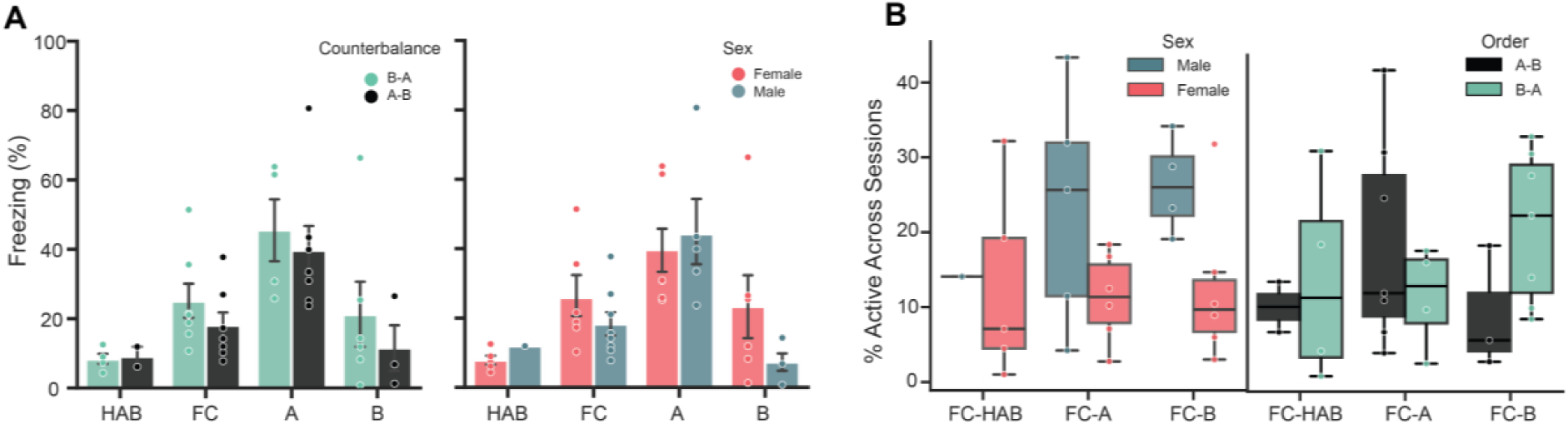
: Behavioral freezing and astrocyte session-level engagement across task phases, stratified by counterbalancing order and sex. **(A)** Freezing behavior (percent time freezing) across task phases (HAB, FC, A, B). Left: data grouped by counterbalancing order (A→B vs B→A). Right: data grouped by sex (female vs male). Bars show group means with error bars indicating uncertainty (SEM), and points denote individual subjects. Linear mixed effects modeling (fixed effects: Sex, Group, Session; random intercept: Mouse) revealed a significant main effect of Session, with increased freezing during FC and Cxt A relative to HAB. No significant main effect of Sex or Group and nor interaction were observed. **(B)** Astrocyte recruitment across paired sessions, quantified as the fraction of astrocytes detected in both sessions and normalized to FC. Distributions are shown as boxplots with overlaid individual subjects, stratified by sex (left) and counterbalancing order (right). Data analyzed using two-way ANOVA on logit-transformed values show that astrocyte recruitment did not differ by counterbalancing order or session pair, and no interaction was observed. In contrast, a significant main effect of sex was detected, and no interaction between sex and session pair. Significance was indicated by: *p ≤ 0.05, **p ≤ 0.01, ***p ≤ 0.001, ****p ≤ 0.0001. See Supplemental Table 1 for statistical details.

**Extended Data Fig. 3.**
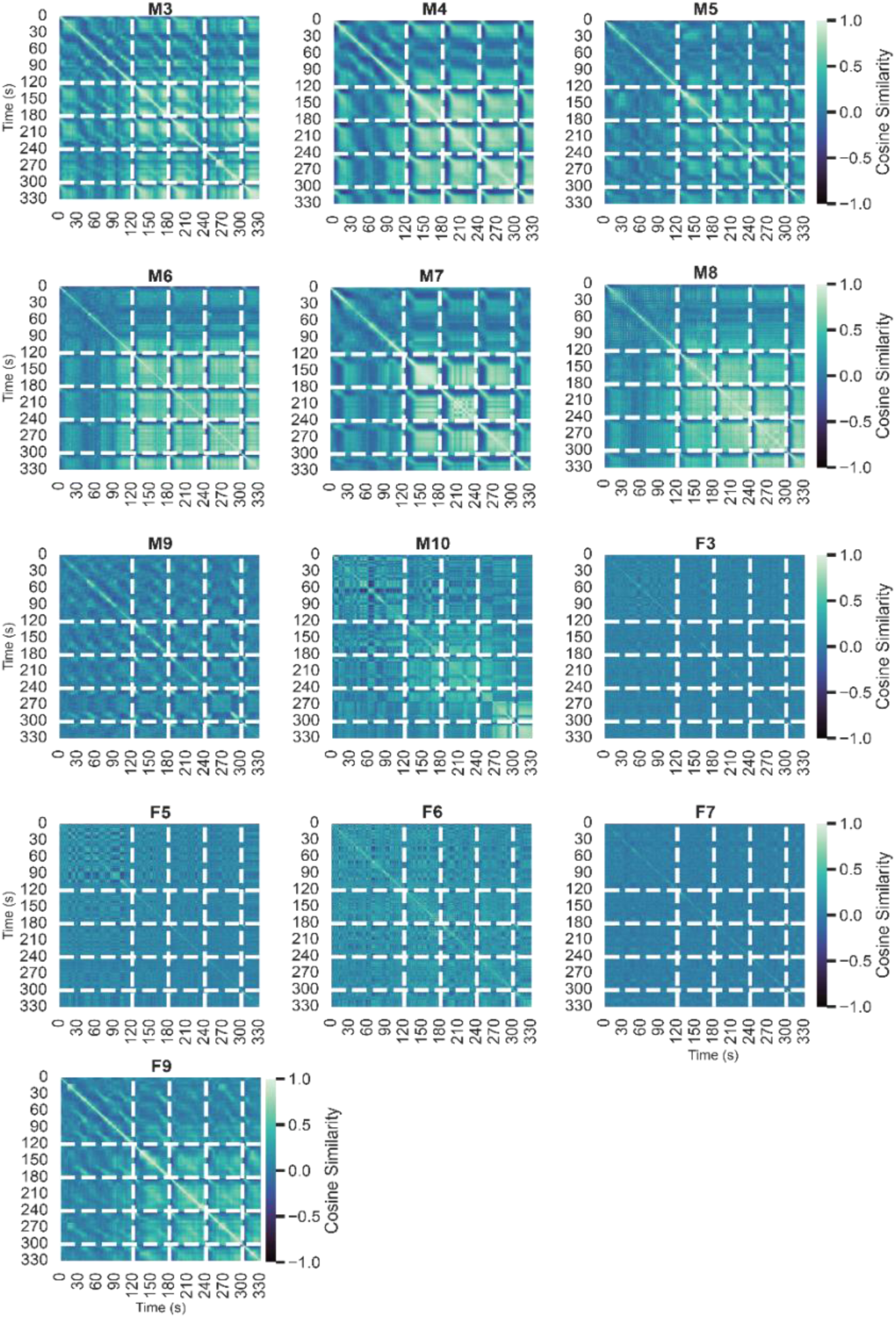
: Individual cosine similarity plots for fear conditioning **(A)** Cosine similarity plots for all animals for contextual fear conditioning, showing the structured temporal dynamics induced by repeated foot shocks. Dashed lines indicate the times of administered foot shocks (120, 180, 240, 300 seconds).

**Extended Data Fig. 4.**
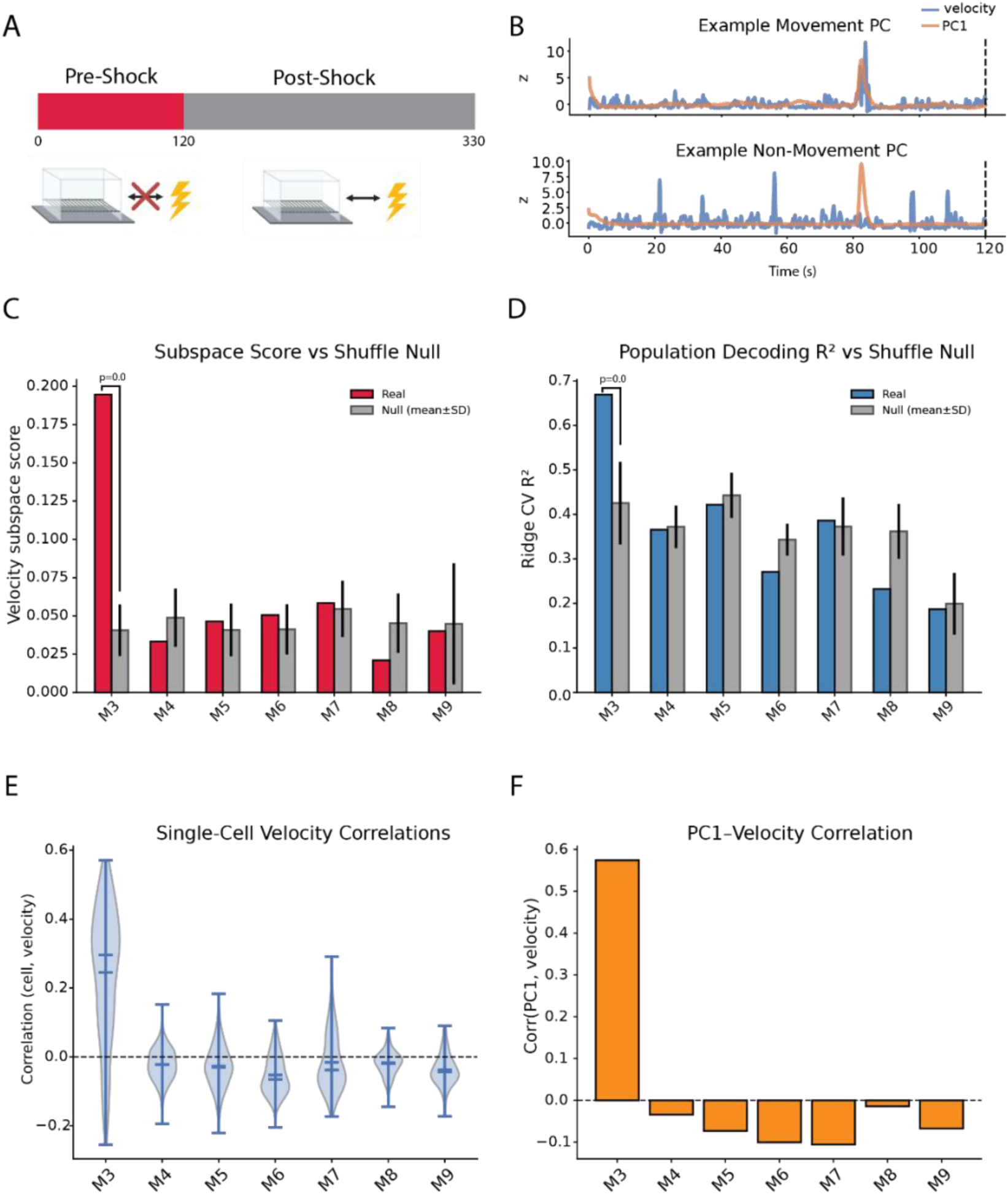
: Relationship between locomotor velocity and astrocytic calcium activity. **(A)** Experimental timeline. Animals are in the pre-shock epoch (0–120 s) until delivery of the first foot shock at 120 s. All behavior here is analyzed in the pre-shock epoch. **(B)** Example principal components (PCs) of astrocytic calcium activity illustrating a movement aligned component (top) and a non-movement component (bottom). Blue traces show animal velocity; orange traces show the corresponding PC time course (PC1). The dashed line marks the transition to the post-shock period. **(C)** Velocity–calcium subspace alignment (“subspace score”) computed for each animal. Colored bars indicate the observed score; gray bars indicate the shuffle-based null distribution (mean ± SD). Only one animal (M3) shows a significant subspace score relative to the null. **(D)** Population decoding of velocity from astrocytic calcium activity using ridge regression. Colored bars show observed cross-validated decoding performance (R²); gray bars show the shuffle null (mean ± SD). Only one animal (astroM3) exhibits significant decoding above the null. **(E)** Distributions of single-cell correlations between each astrocyte calcium trace and concurrently measured velocity for each animal (violin plots). **(F)** Correlation between the dominant calcium PC (PC1) and velocity for each animal; only one animal shows a large positive PC1–velocity correlation. Unless noted otherwise, error bars denote mean ± SEM; for shuffle controls, gray bars denote mean ± SD. Significance: *p ≤ 0.05, **p ≤ 0.01, ***p ≤ 0.001, ****p ≤ 0.0001. See Supplemental Table 1 for statistical details.

**Extended Data Fig. 5.**
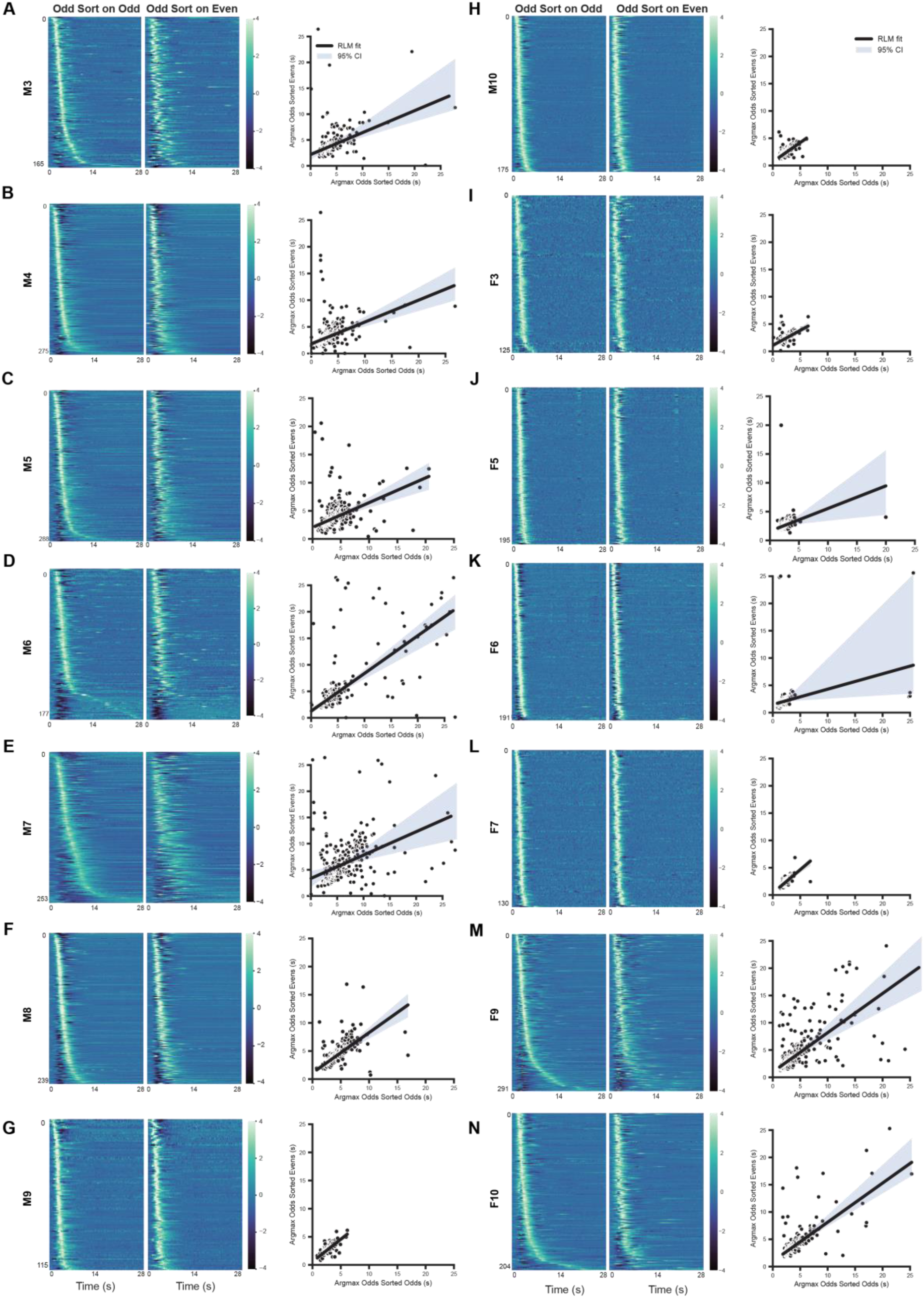
: Cross validated heatmaps for contextual fear conditioning. **(A-N; left)** Heatmaps of all astrocytic calcium activity that was averaged across shocks #1 and 3 (Odds) and sorted by their peak amplitude during FC. To assess the stability of the sequential response, the same activity for Odds was sorted based on the peak amplitude of the average response of each astrocyte to shocks #2 and 4 (Evens). Here, we observe a stable sequence across all animals across foot shocks (n = 14). Spearman’s correlation calculated between the true peak times (Odd sorted on Odd) and cross-validated peak times (Odd sorted on Even) showed a significant difference between the average rho value and shuffle null using a paired t-test (Figure 2e). **(A-N; right)** Linear regressions of the true peak times (i.e., Odd Sort on Odd Peaks) against the cross-validated peak times (i.e., the Odd Peaks sorted on the Even Peak times) for all animals. The solid line indicates the fixed effects best fit line that predicts the even peak-times from the odd-peak times with the associated 95% CI on each plot. See Supplemental Table 1 for statistical details.

**Extended Data Fig. 6.**
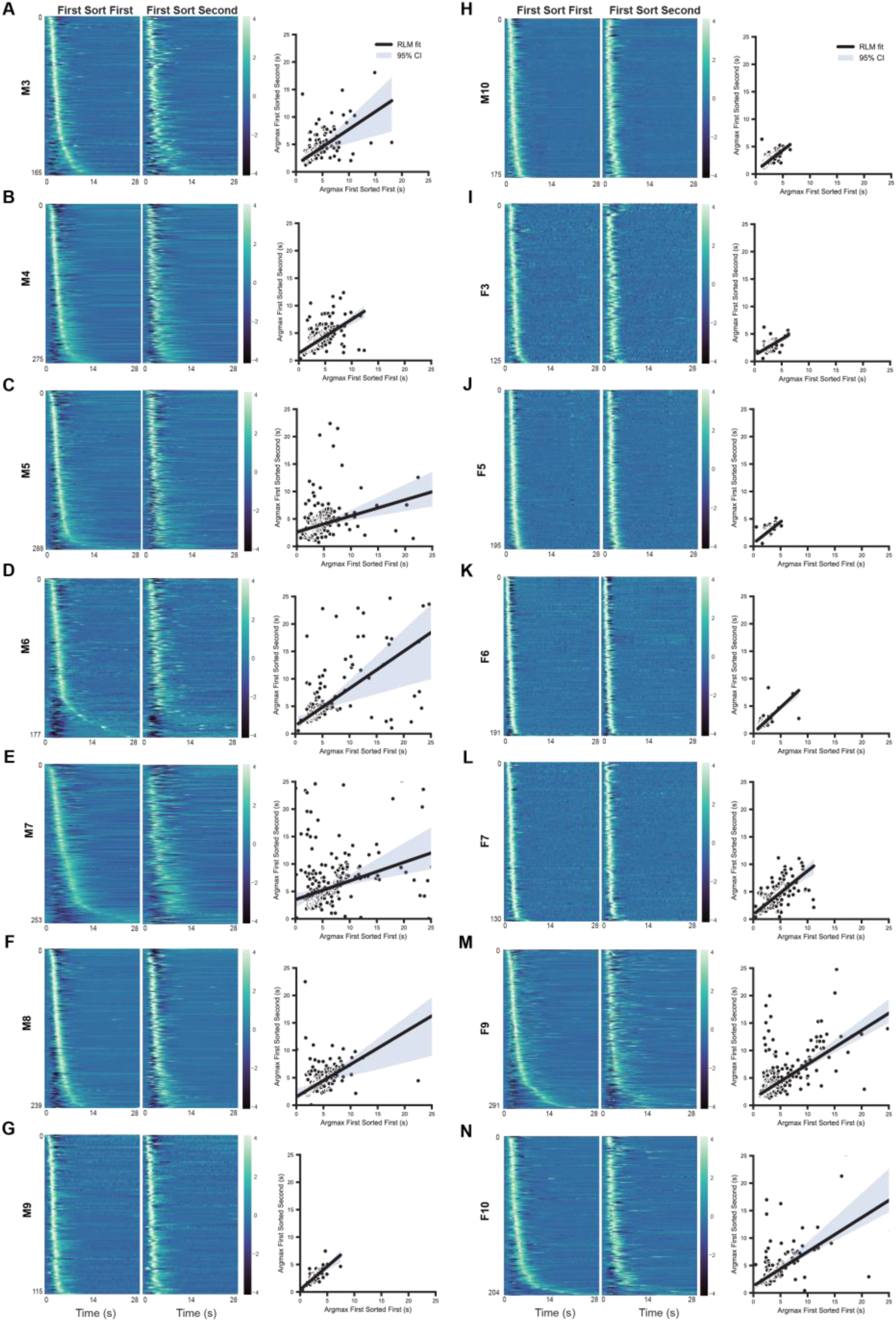
: Cross-validation of first and second half of fear conditioning plots **(A–N; left)** Heatmaps of astrocytic calcium activity aligned to foot shocks, averaged separately across the first half of shocks (Early) and the second half of shocks (Late). Traces are shown as z-scored activity (or normalized activity, as in the main analysis) for all astrocytes, and heatmaps are sorted by each astrocyte peak response time during the Early average. To assess the stability of the sequential response from early to late trials, the same Early activity is re-ordered using the peak response times computed from the Late average (cross-validated sorting). We observe a stable response sequence across animals from early to late shocks (n = 14). Stability was quantified using Spearman’s correlation between true peak times (Early sorted on Early peaks) and cross-validated peak times (Early activity sorted using Late-derived peak times). The mean Spearman’s ρ across animals was compared against a shuffle-null distribution (paired t-test: t=18.788, p<0.0001). **(A–N; right)** Linear regressions relating true peak times (Early sort on Early peaks) to cross-validated peak times (Early peaks ordered by Late-derived peak times) for each animal. The solid line shows the fixed-effects best-fit relationship predicting the even peak-times from the odd peak-times, with the associated 95% confidence interval. See Supplemental Table 1 for statistical details.

**Extended Data Fig. 7.**
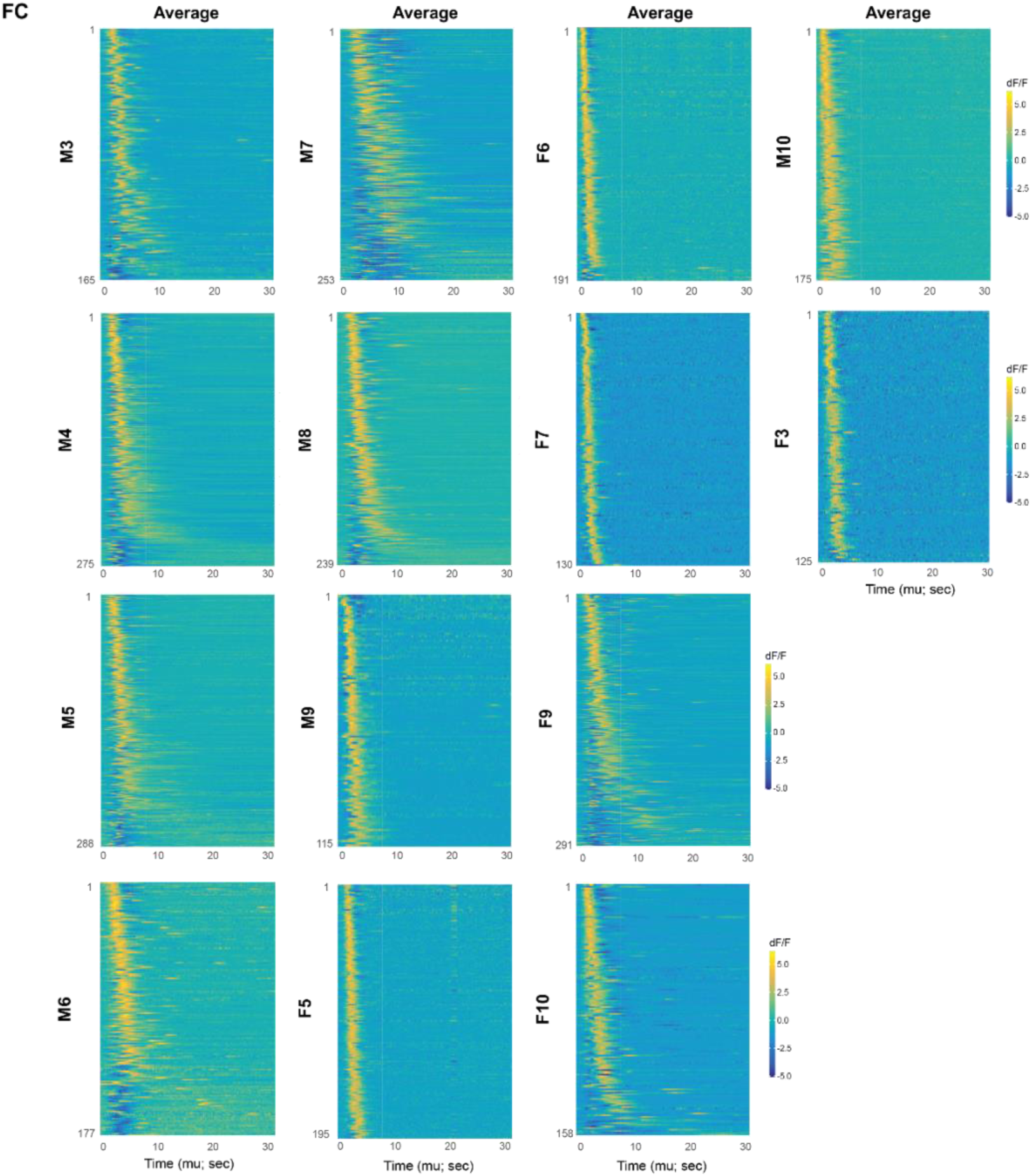
: Individual shock heatmaps for all animals derived from Bayesian modeling for contextual fear conditioning. Average heatmaps of individual astrocytic calcium traces in response to foot shock for all mice. These were sorted based on the average predicted peak times (µ) that were extracted from the Hierarchical Bayesian Model. Lower values of mu (i.e., cells that had earlier peak times) are plotted on the top (i.e., ascending order). Average (left) and individual detected sequences (middle). See Methods for model specifications and details.

**Extended Data Fig. 8.**
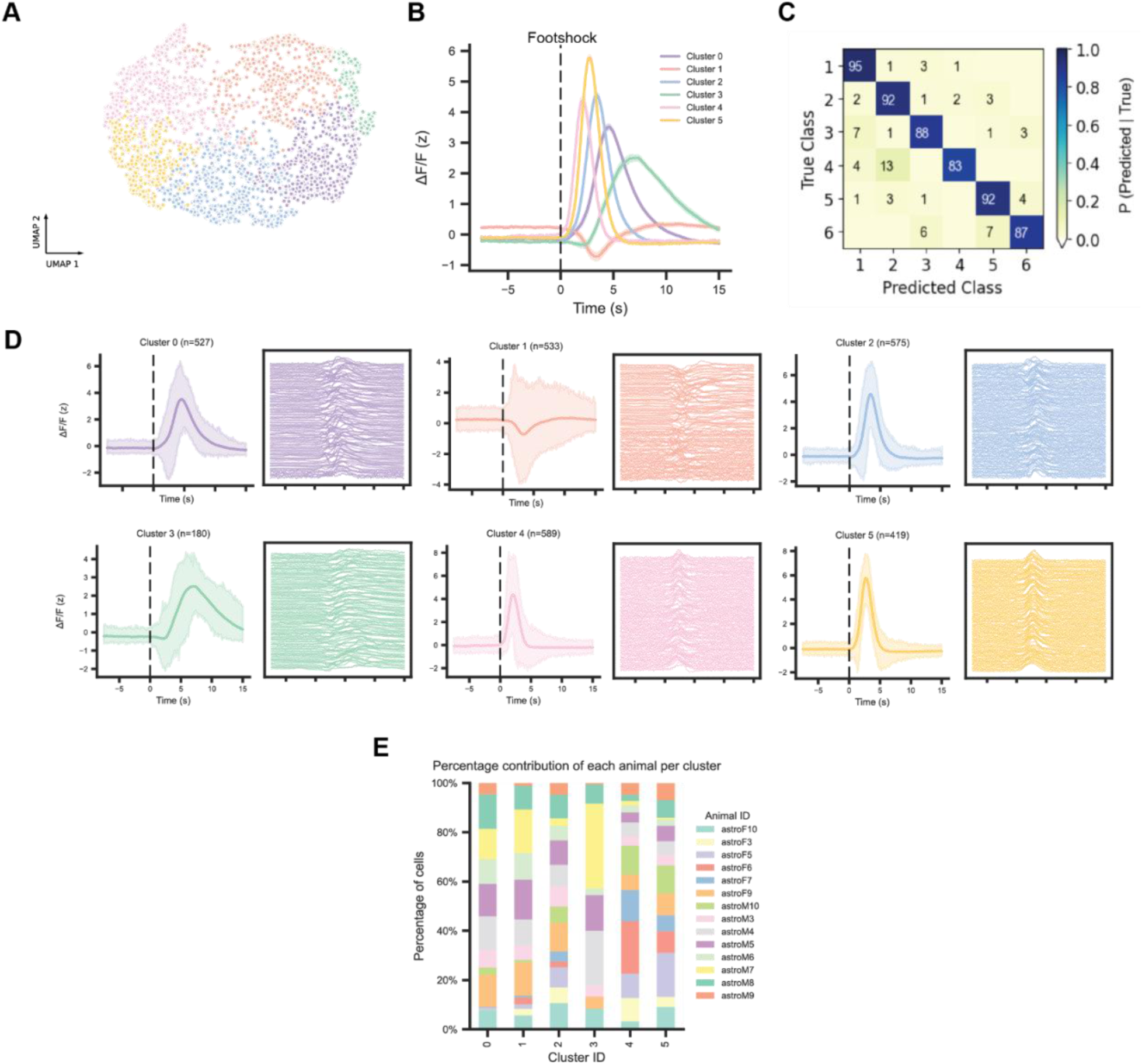
:Exploratory investigation of astrocytic calcium dynamics in response to foot shock using WaveMAP. **(A)** WaveMAP results for all mice during FC represented in two-dimensional UMAP space. The colormap is matched across Extended Data Fig. 2A-D to represent each cluster. **(B)** Average foot shock response (z-scored % dF/F) for each of the 6 clusters (cluster 0-5) detected by WaveMAP, displaying unique kinetic characteristics that vary with amplitude and decay time. Shaded regions indicate the t-confidence interval for each cluster. **(C)** Confusion matrix displaying high accuracy in correctly classifying calcium waveforms within their respective clusters. These matrices were used to optimize WaveMAP parameters that would most accurately cluster astrocytic waveforms (n_neighbors=30; mindistance=0.1; n=2823 cells). **(D)** (Left panels) Average calcium waveforms (z-scored % dF/F) within each detected cluster at the onset of foot shock (dashed line; t = 0) (0-5). n values represent the number of astrocytes that fall within that cluster. Shaded regions indicate the bootstrapped confidence interval for each cluster. (Right panels) Individual calcium waveforms for n = 100 single-astrocytes within each detected cluster to show slight variation of dynamics within the cluster. **(E)** Percentage of astrocytes for each animal represented within a respective cluster (0-5) show that clusters are not driven by individual animal differences. Colors here do not correspond to the cluster colors and instead correspond with an animal. See Supplemental Table 1 and Methods for statistical details.

**Extended Data Fig. 9.**
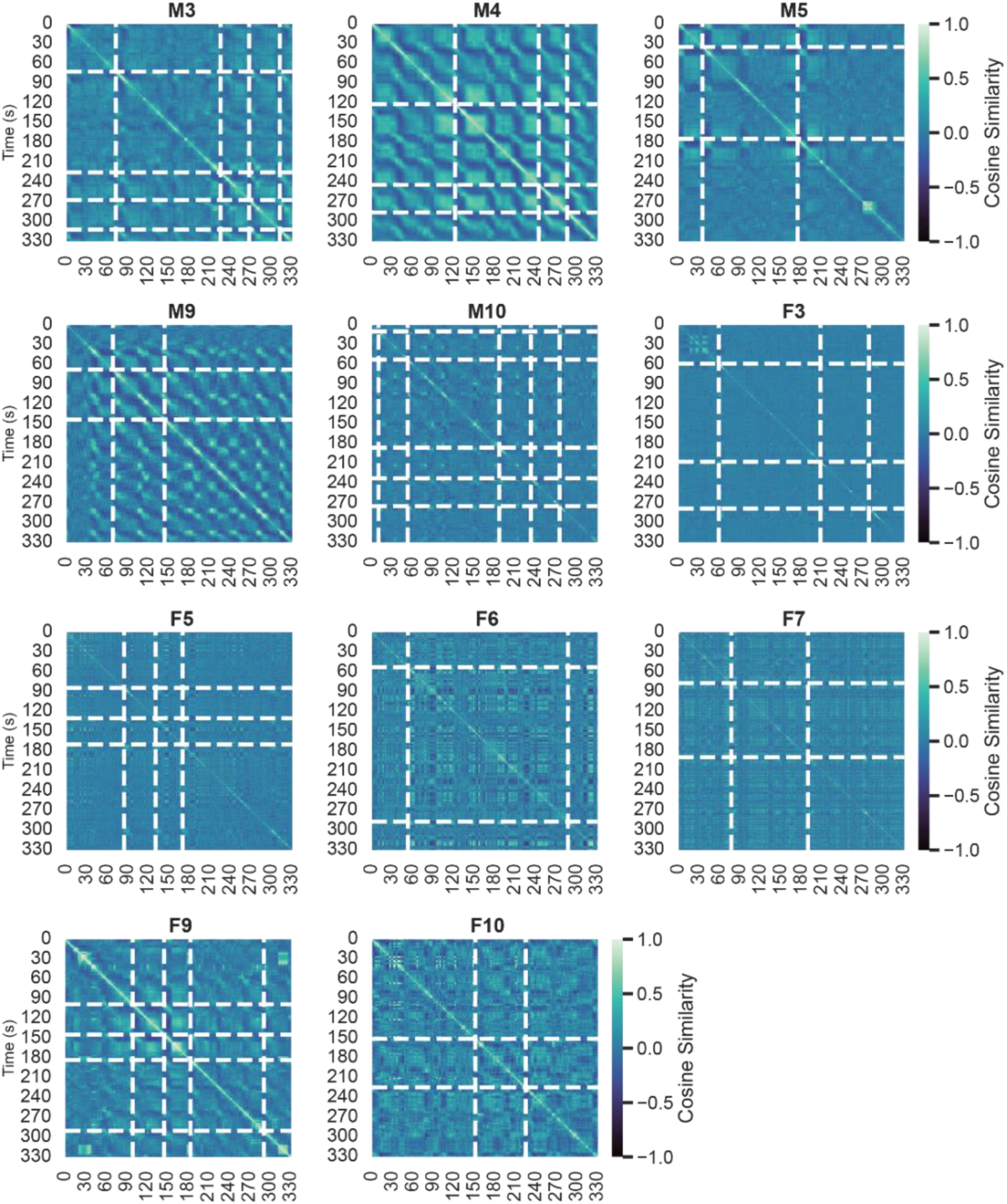
: Individual cosine similarity plots for context A Cosine similarity plots for all animals for re-exposure to context A. White lines indicate detected high activity “events” from the sequence detector.

**Extended Data Fig. 10.**
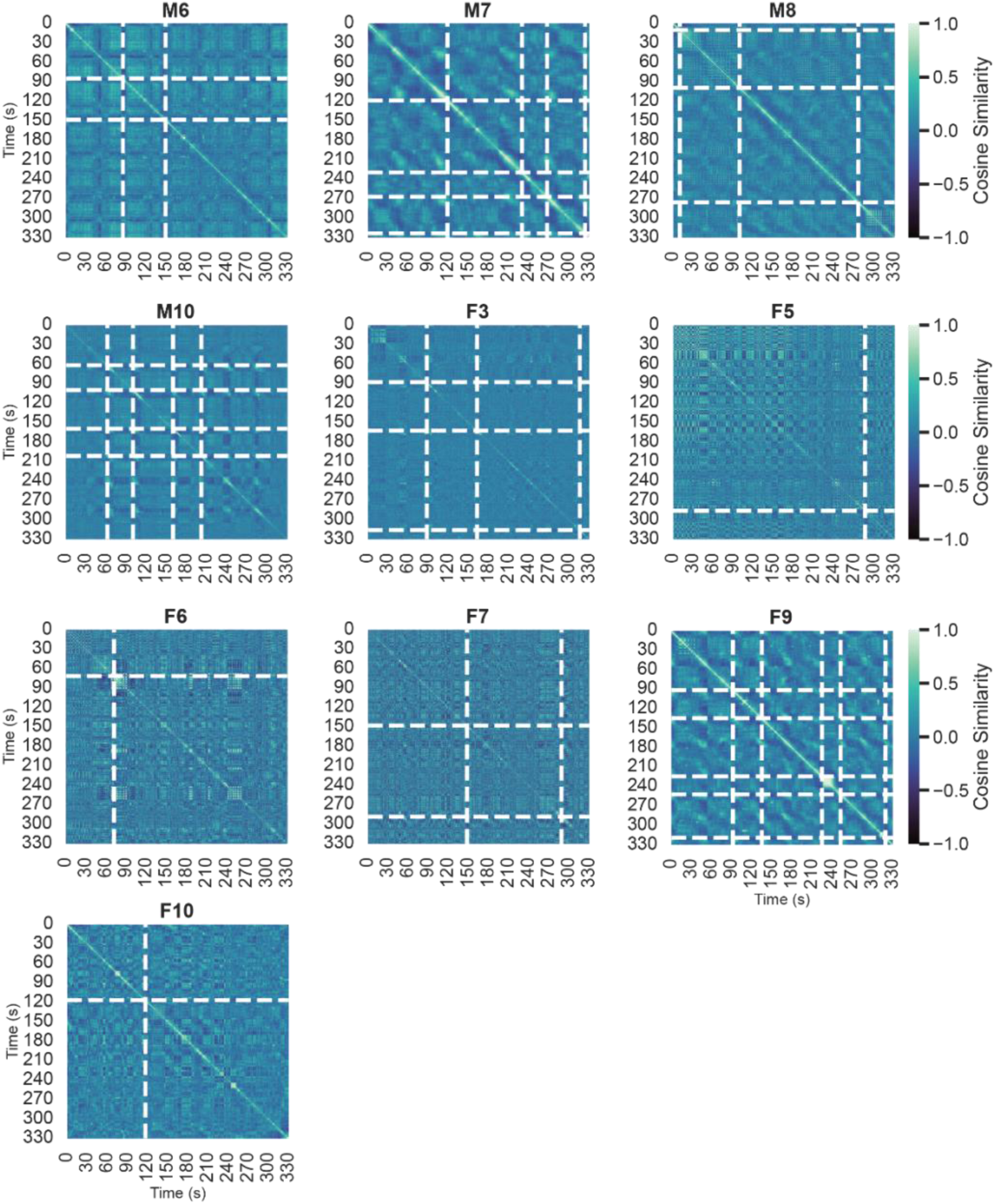
: Individual cosine similarity plots for context B Cosine similarity plots for all animals for novel exposure to context B. White lines indicate detected high activity “events” from the sequence detector.

**Extended Data Fig. 11.**
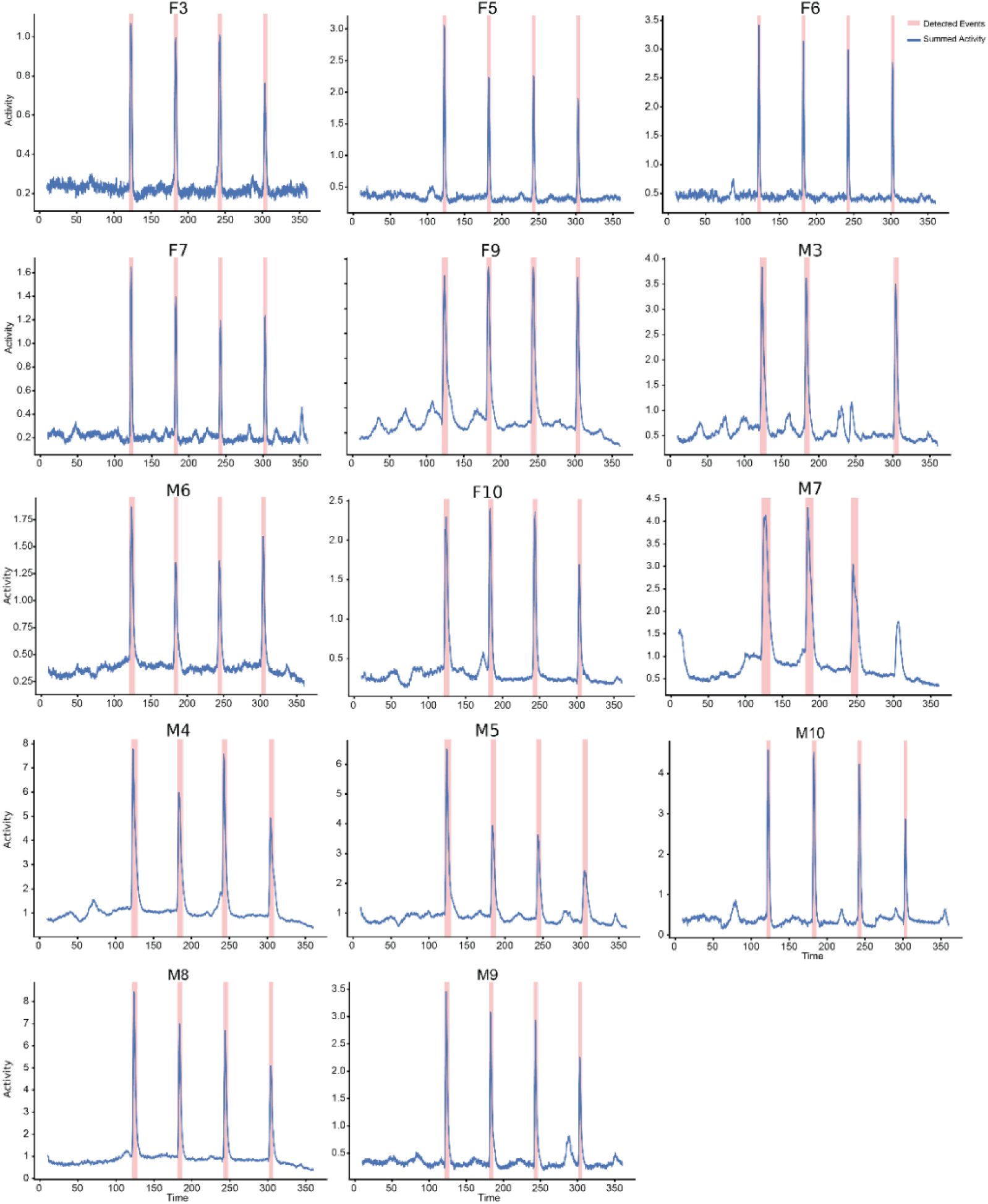
: Individual animal high-activity event detector results for contextual fear conditioning. Detected shock sequences for all individual animals during contextual fear conditioning as a confirmation of the ‘sequence detector’ described in Fig. 3d. Individual calcium activity was summed over each time index to generate the summed activity trace for a given mouse (solid line; Cxt A = blue, Cxt B = black). Temporal thresholding was performed by finding the time indices where the summed activity was > 1.0 standard deviations above the mean and lasted for > 2 seconds. These time periods qualify as detected ‘sequences’ (shading). Please note that two mice M3 and M7 only detected n=3 shock events, as the mice were able to shift their body partially off the grid to minimize the amount of shock received during that trial (i.e. lower amplitude).

**Extended Data Fig. 12.**
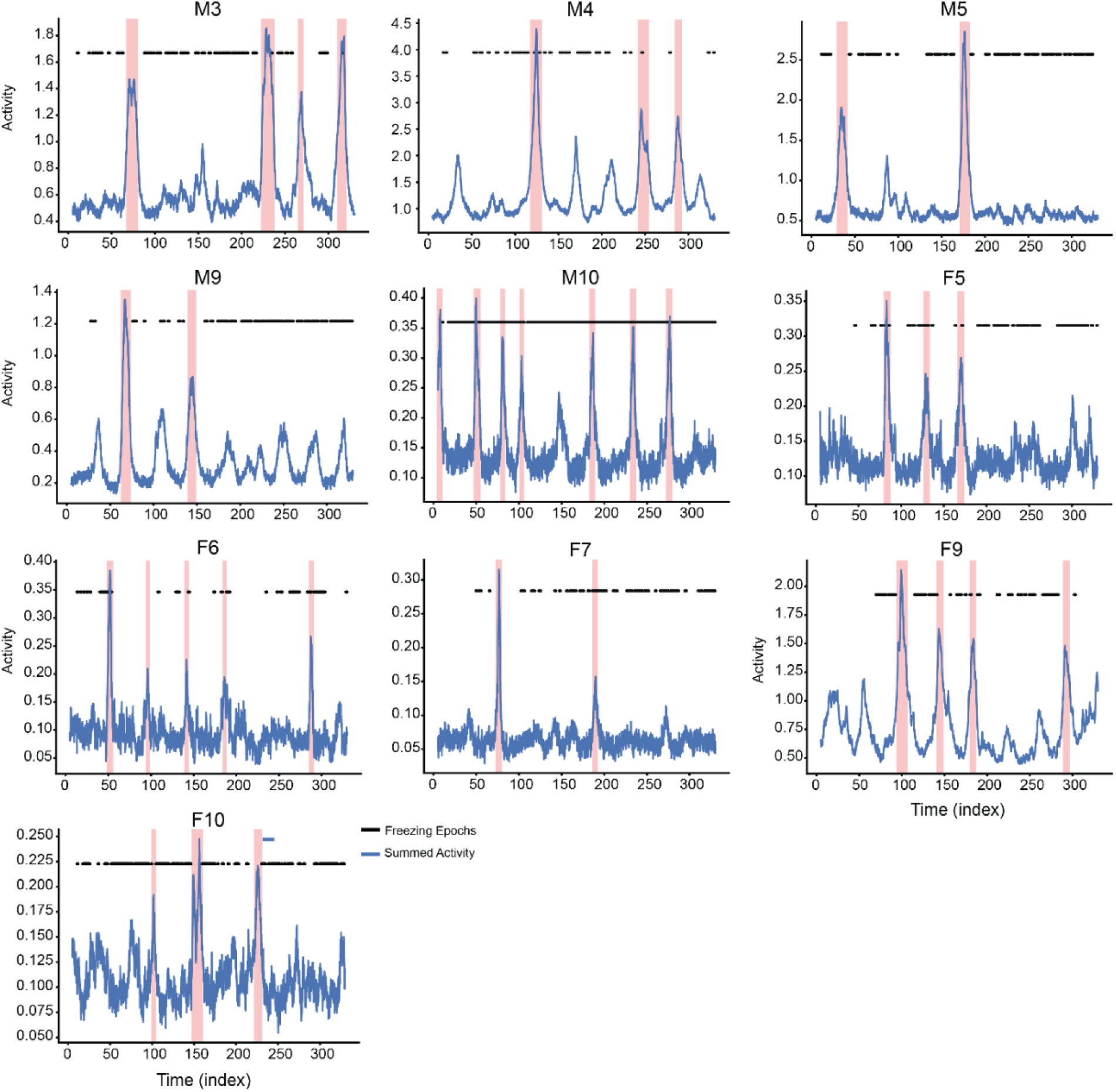
: Individual animal high-activity event detector results for context A recall. Detected high activity sequences for all individual animals in Cxt A described in Fig. 3. Individual calcium activity was summed over each time index to generate the summed activity trace for a given mouse (solid line; blue). Temporal thresholding was performed by finding the time indices where the summed activity was > 1.0 standard deviations above the mean and lasted for > 2 second. These time periods qualify as detected ‘sequences’ (shading). Freezing periods are shown as black dashes across the session.

**Extended Data Fig. 13.**
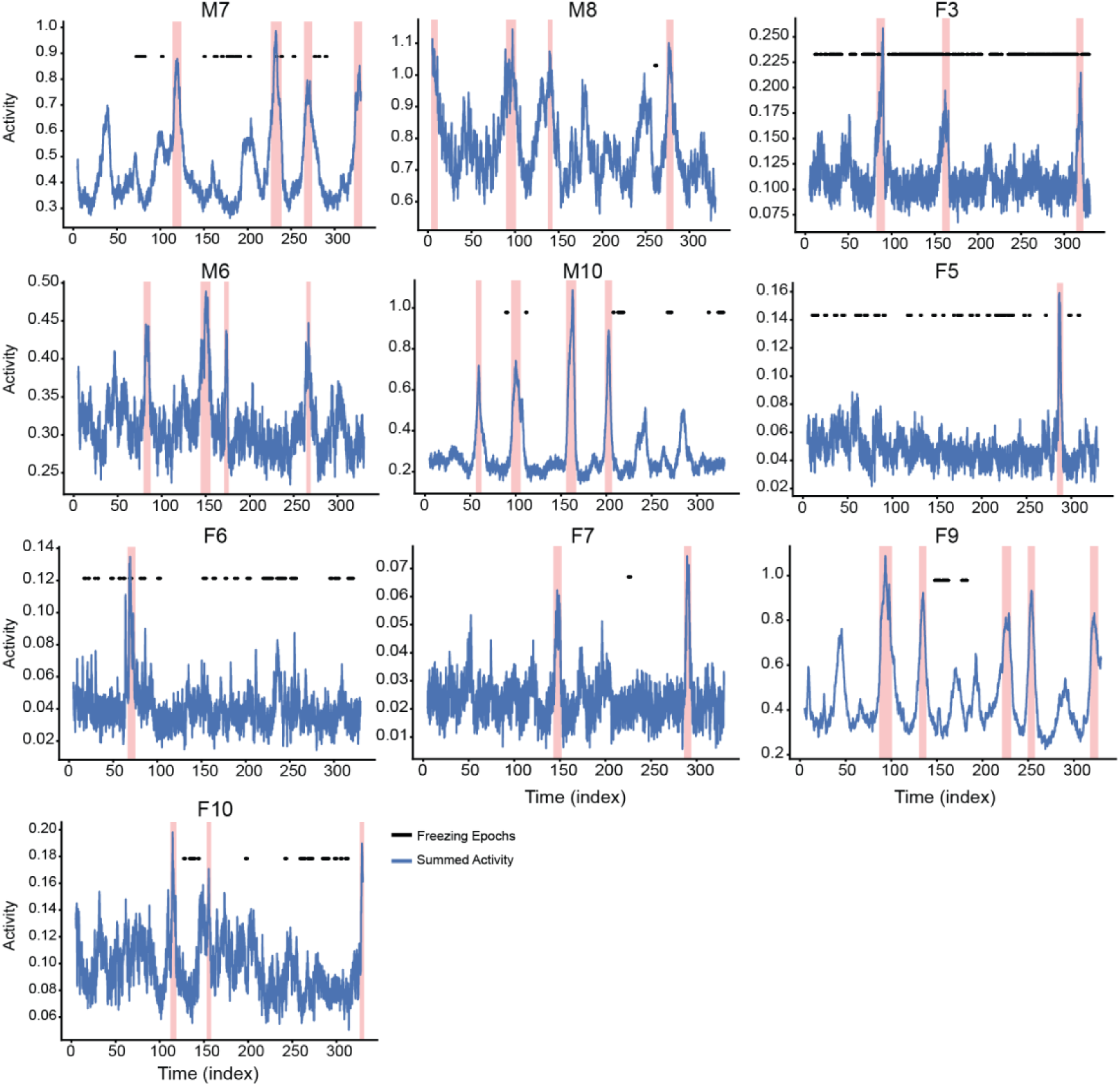
: Individual animal high-activity event detector results for context B Detected high activity sequences for all individual animals in Cxt B described in Fig. 3. Individual calcium activity was summed over each time index to generate the summed activity trace for a given mouse (solid line; blue). Temporal thresholding was performed by finding the time indices where the summed activity was > 1.0 standard deviations above the mean and lasted for > 2.0 second. These time periods qualify as detected ‘sequences’ (shading). Freezing periods are shown as black dashes across the session.

**Extended Data Fig. 14.**
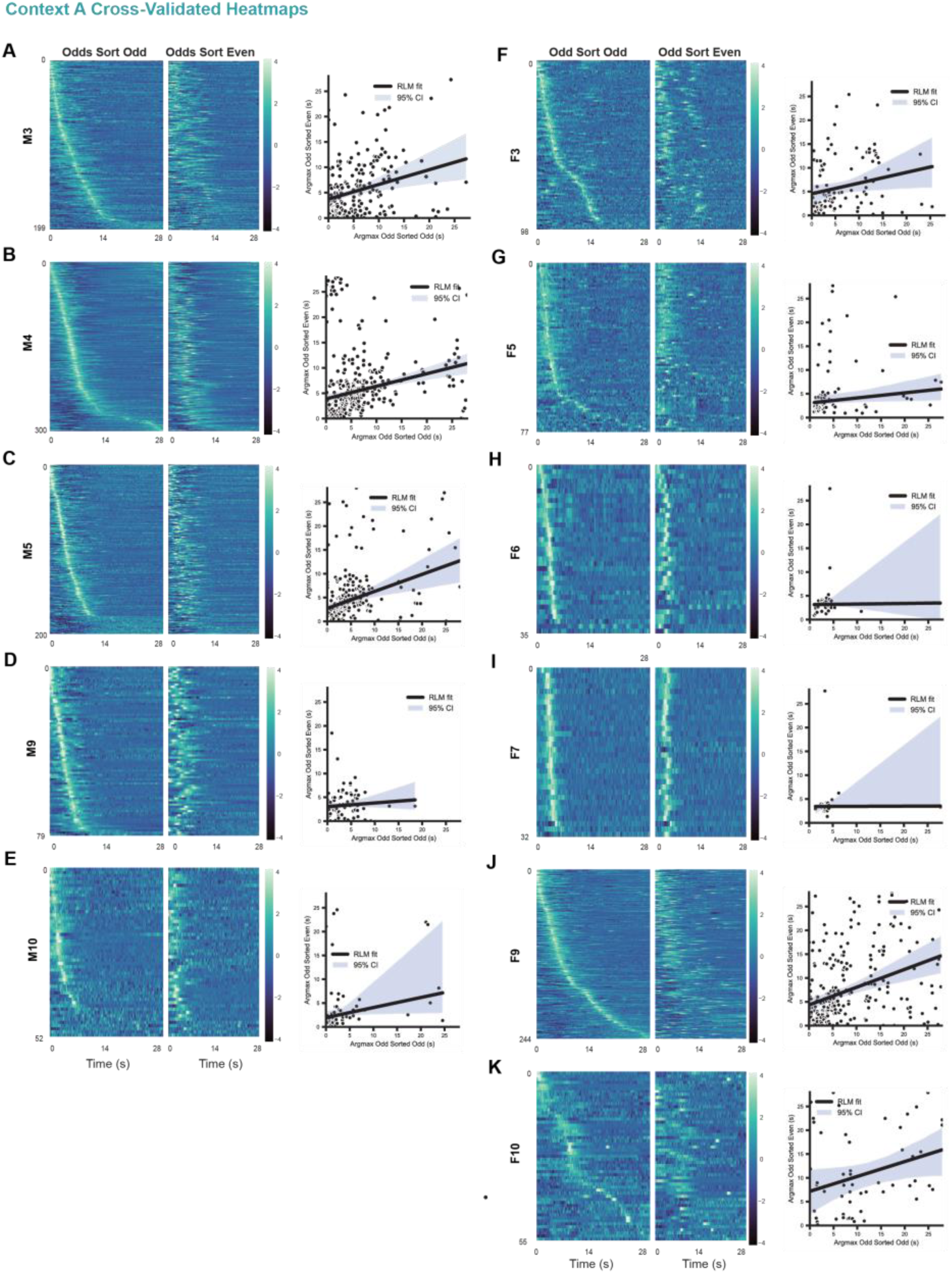
: Cross validated heatmaps for context A **(A-K; left)** Heatmaps of all astrocytic calcium activity that were aligned to detected events (Odds) and sorted by their peak amplitude (Odds Sort Odds) for exposure to Cxt A. To assess the stability of the sequential response, the same activity for Odds was sorted based on the peak amplitude of the average response of each astrocyte to spontaneously detected events (Evens). **(A-K; right)** Linear regressions of the true peak times (i.e., Odd Sort on Odd Peaks) against the cross-validated peak times (i.e., the Odd Peaks sorted on the Even Peak times) for all animals. The solid line shows the robust linear model best-fit relationship predicting the even peak-times from the odd peak-times, with the associated 95% confidence interval. See Supplemental Table 1 for full model specifications and statistics. Confidence intervals are bootstrapped at a 95% level.

**Extended Data Fig. 15.**
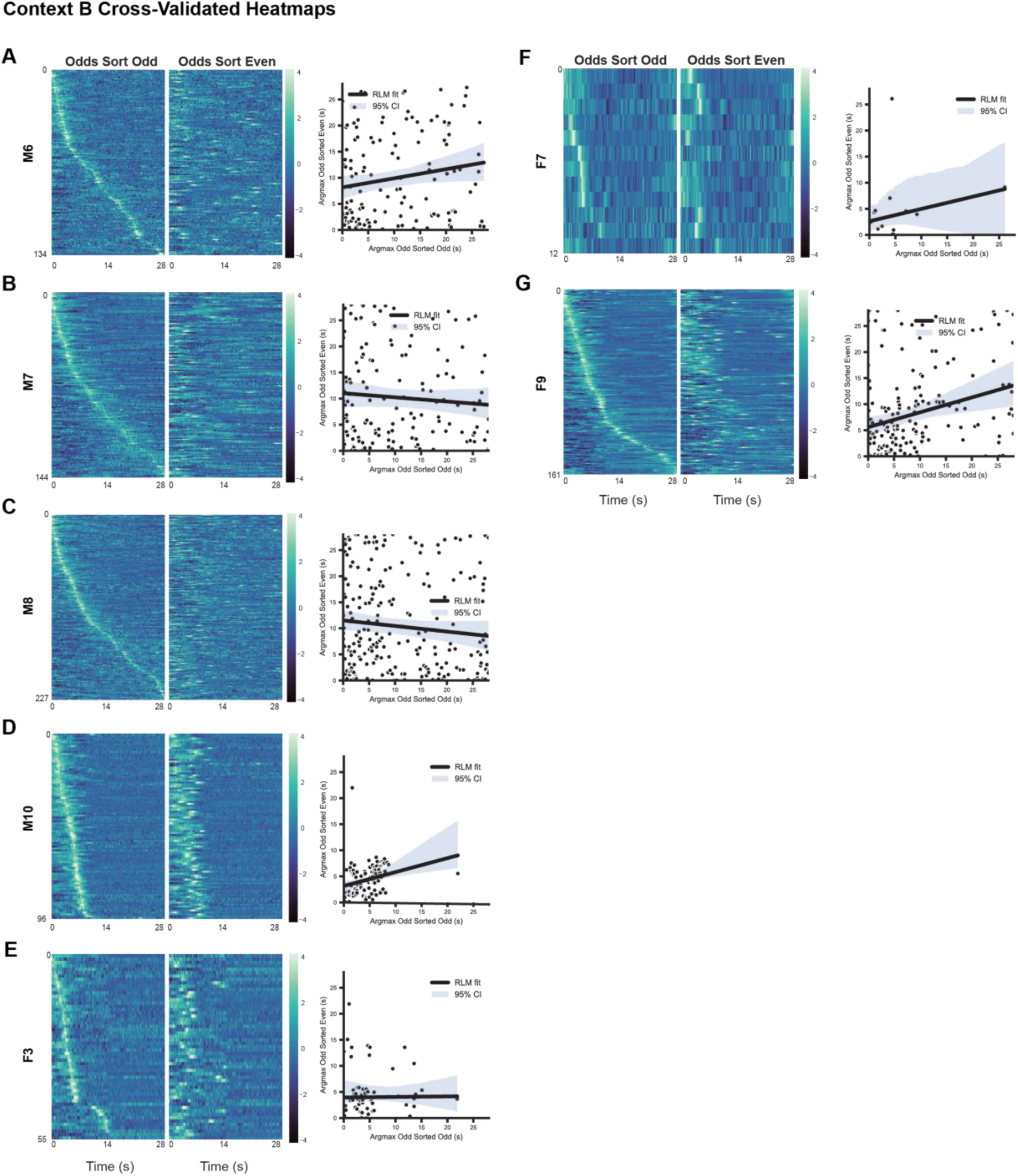
: Cross validated heatmaps for context B **(A-G; left)** Heatmaps of all astrocytic calcium activity that were aligned to detected events (Odds) and sorted by their peak amplitude (Odds Sort Odds) for exposure to Cxt B. To assess the stability of the sequential response, the same activity for Odds was sorted based on the peak amplitude of the average response of each astrocyte to spontaneously detected events (Evens). **(A-G; right)** Linear regressions of the true peak times (i.e., Odd Sort on Odd Peaks) against the cross-validated peak times (i.e., the Odd Peaks sorted on the Even Peak times) for all animals. The solid line shows the robust linear model best-fit relationship predicting the even peak-times from the odd peak-times, with the associated 95% confidence interval. See Supplemental Table 1 for full model specifications and statistics. Confidence intervals are bootstrapped at a 95% level.

**Extended Data Fig. 16.**
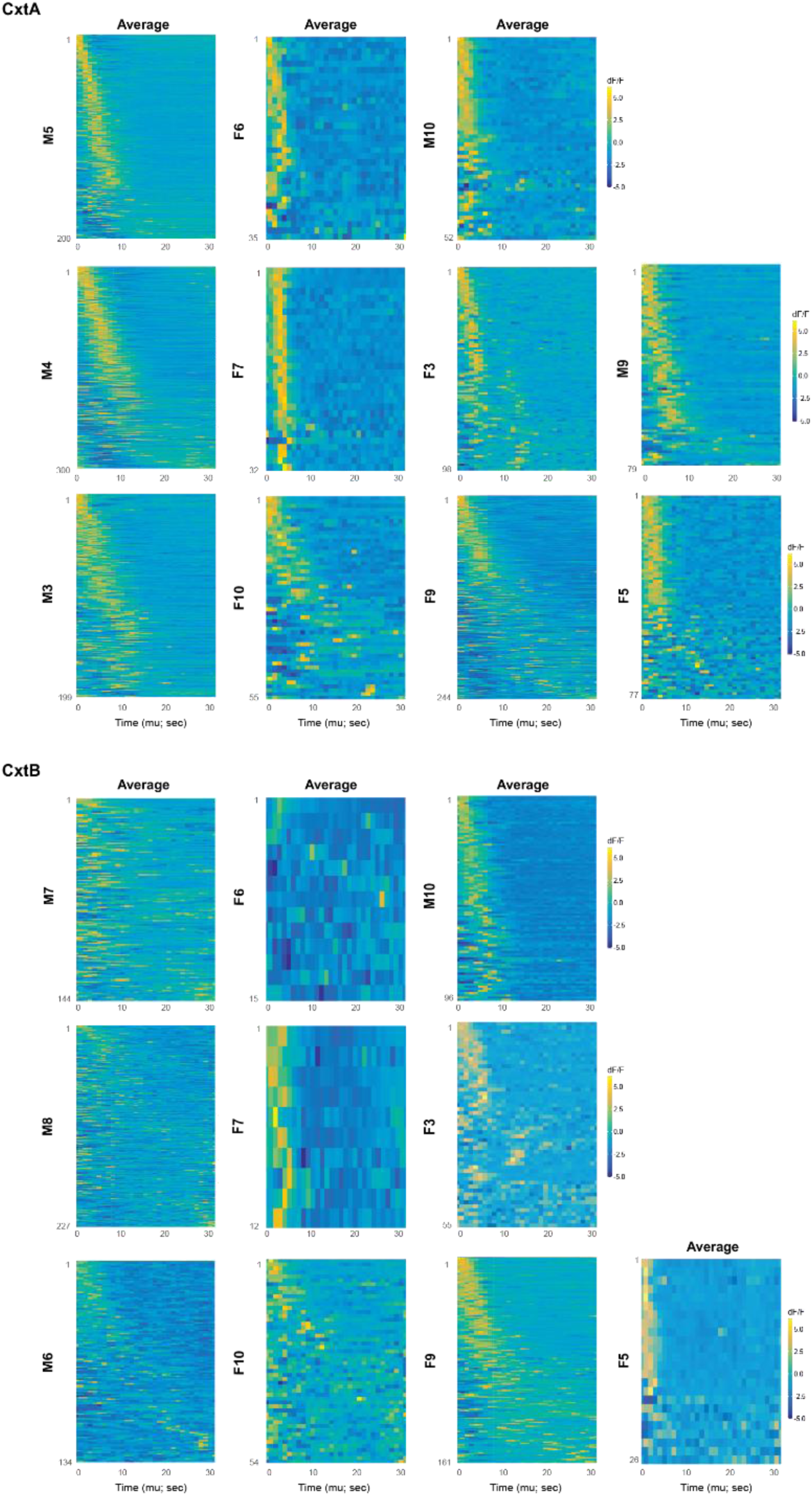
: Individual heatmaps for all animals derived from Bayesian modeling for recall. *Top.* Analysis for Cxt A. Average heatmaps of individual astrocytic calcium traces in response to foot shock for all mice. These were sorted based on the average predicted peak times (µ) that were extracted from the Hierarchical Bayesian Model. Lower values of mu (i.e., cells that had earlier peak times) are plotted on the top (i.e., ascending order). Average (left) and individual detected sequences (middle). See Methods for model specifications and details. *Bottom.* Same as above but for exposure to Cxt B.

**Extended Data Fig. 17.**
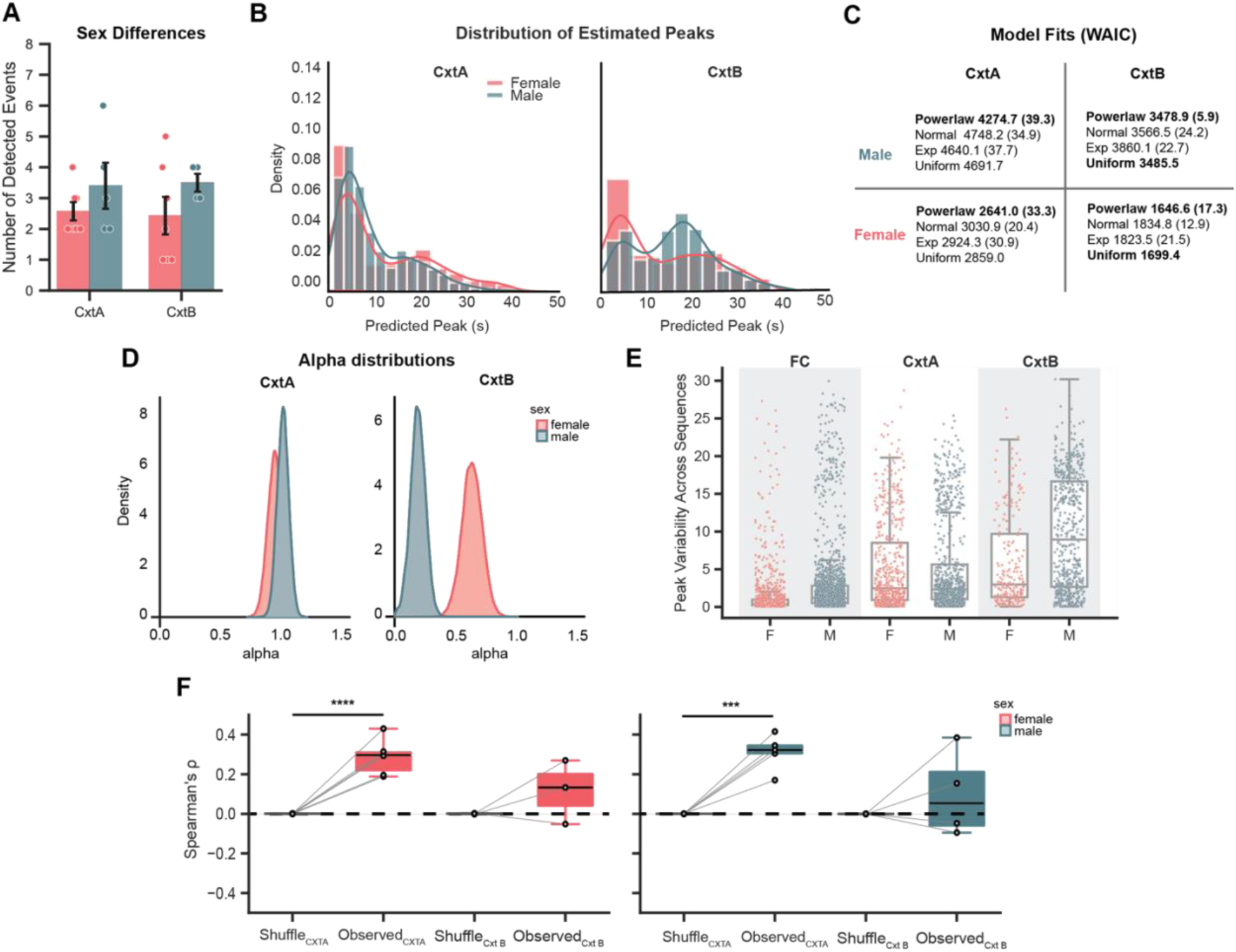
: Analysis of potential sex differences in event detection and model-based characterization of peak structure across contexts. **(A)** Mean number of detected events in two contexts (Cxt A, Cxt B) shown separately for females (red) and males (blue). Points denote individual subjects; bars indicate group means with error bars reflecting uncertainty (SEM). There were no significant differences in the number of detected events across sex or context (Two-way ANOVA; Context and Sex as main effects). **(B)** Group-level distributions of estimated peak location (predicted peak, μ) for Cxt A and Cxt B, shown as histograms with smoothed density overlays for females and males. Cxt A (male; n=5, cells=830) (female; n=6, cells=541). Cxt B (male; n=4, cells=601) (female; n=6, cells=323). **(C)** Model comparison (WAIC) for candidate distributions used to describe estimated peak locations in each context and sex group (Power-law, normal, exponential, uniform). Lower WAIC indicates better expected out-of-sample predictive performance; values in parentheses report the associated uncertainty. Best-supported model(s) are highlighted in bold. **(D)** Posterior distributions of the Power-law scaling parameter (α) by context and sex. In context B, posterior distributions are non-overlapping suggesting clear peak distributional differences, while in context A, these posterior distributions are largely overlapping. **(E)** Peak variability across sequences (FC) and detected high activity event periods for Cxt A and Cxt B, summarized by boxplots with overlaid individual astrocyte observations (F, female; M, male). Linear mixed effects modeling showed a main effect of Session (FC vs Cxt A, p<0.0001) and significant Sex x Session interactions for both Cxt B and FC (p<0.0001). This shows that males exhibit increased variability in Cxt B and a reduced FC-related decrease in peak variability compared to females. Cxt A (male; n=819 cells) (female; n=535 cells); Cxt B (male; n=596 cells) (female; n=227 cells); FC (male; n=1672 cells) (female; n=1074 cells). **(F)** Within-subject Spearman rank correlations (ρ) quantifying consistency in peak structure across sequences, comparing observed data to a shuffled control for each Context and Sex. Both sexes in Cxt A were significantly more correlated than the shuffle null distribution (paired t-test [female: t=7.87, p<0.001] [male: t=7.72, p=0.002]), while both sexes in Cxt B were not (paired t-test: [female: t=1.24, p=0.340] [male: t=0.90, p=0.432]). Lines connect paired shuffled vs observed estimates within subject. Mice with < 2 detected events were removed from this analysis (Cxt A: male; n=5, female; n=6) (Cxt B: male; n=4, female; n=3). Significance annotations indicate inferential results (*p ≤ 0.05, **p ≤ 0.01, ***p ≤ 0.001, ****p ≤ 0.0001, ns). See Supplemental Table 1 for statistical details.

**Extended Data Fig 18.**
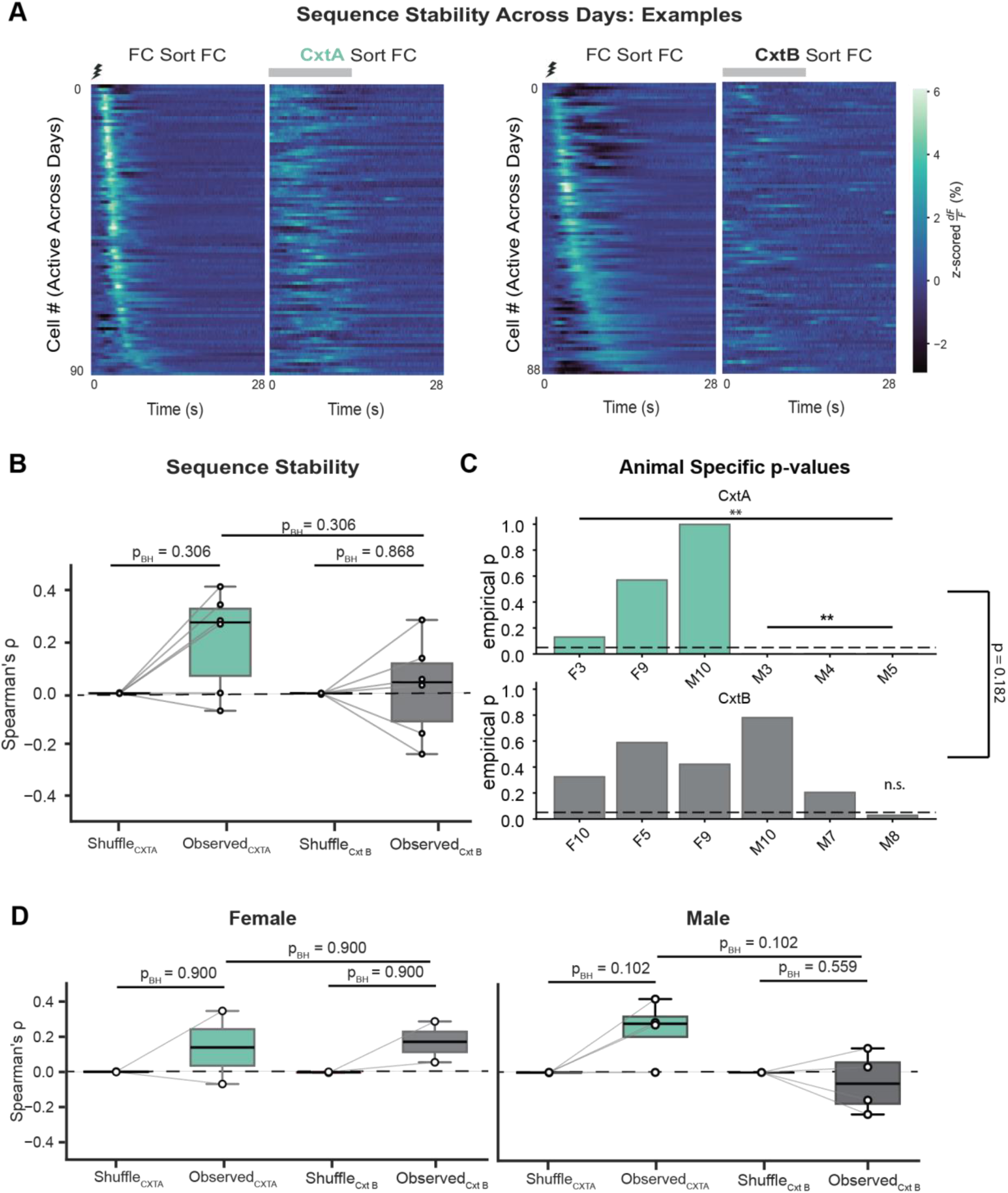
: Cross-day astrocyte sequence order showed variable preservation across animals. (A) Example heatmaps of calcium activity from astrocytes active across days during fear conditioning (FC) and recall in Context A (left) or Context B (right). Astrocytes were sorted by peak response time during FC, and this same ordering was then applied to activity during recall in Context A or Context B. Activity is shown as z-scored dF/F (%) aligned to foot shock during FC (shock symbol) or to detected spontaneous events during recall (grey bar). (B) Spearman rank correlations comparing astrocytic peak times during FC with cross-validated peak times during recall for cells active across both days (Context A, blue, n = 6; Context B, black, n = 6). At the group level, neither context differed significantly from its shuffle-based null distribution (paired t-tests: Cxt A, p = 0.140; Cxt B, p = 0.868). Null-corrected correlations (observed minus shuffle mean) also did not differ between Cxt A and Cxt B (independent t-test: p = 0.204). Animals with fewer than 10 astrocytes active across days were excluded from this analysis (see Extended Data Fig. 19 for individual animals). Points represent individual animals; grey lines connect observed and shuffled values within each context. The dashed horizontal line indicates the approximate shuffle baseline. (C) Individual-animal p-values from a bootstrap hypothesis test of cross-day Spearman’s ρ values. Three of six animals in Cxt A showed significant cross-day sequence correlations after Benjamini–Hochberg correction, whereas none of six animals in Cxt B did. Exact binomial tests indicated that the 3/6 result in Cxt A was significant relative to a null success probability of *p* = 0.05, whereas the 0/6 result in Cxt B was not. However, a Fisher’s exact test did not reject equality of proportions between contexts, indicating that these results are heterogeneous and should be interpreted cautiously. The dashed line indicates the nominal significance threshold of 0.05 before multiple-comparison correction. (D) Spearman rank correlations comparing astrocytic peak times during FC with cross-validated peak times during recall, shown separately by sex (Context A, blue; Context B, black). In females, correlations did not differ from shuffle controls in either context, and no difference was observed between contexts (Context A, n = 2; Context B, n = 2). In males, neither context differed significantly from shuffle after accounting for multiple-hypothesis test correction (Context A, n = 4; Context B, n = 4). Animals with fewer than 10 astrocytes active across days were excluded from this analysis. Points represent individual animals; grey lines connect observed and shuffled values within each context. The dashed horizontal line indicates the approximate shuffle baseline.

**Extended Data Fig. 19.**
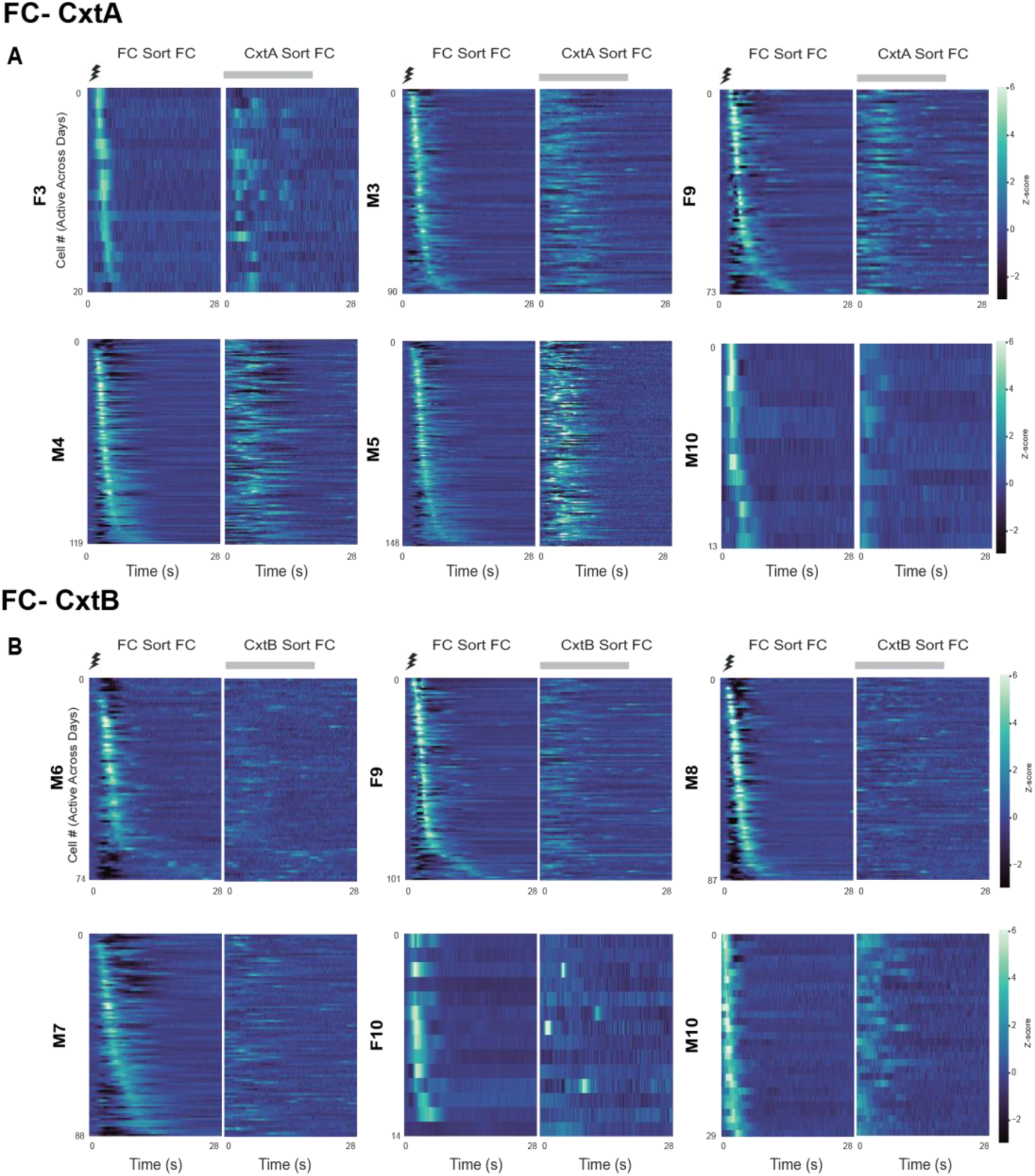
: Cross-validated heatmaps across FC-A and FC-B session pairs using astrocytes active across days. **(A)** FC-Cxt A and **(B)** FC-Cxt B. *Left panel:* Heatmap of individual astrocytes that were active across sessions in response to foot shock during fear conditioning (FC) sorted on their peak amplitude. *Right heatmaps*: Heatmap of the same astrocytes averaged over spontaneously detected events, but during re-exposure to Cxt A or exposure to Cxt B. Heatmap is sorted on the peak amplitude values from FC the day prior. Mice that were removed from analysis (n < 10 cells across days) were not shown here.

**Extended Data Fig. 20.**
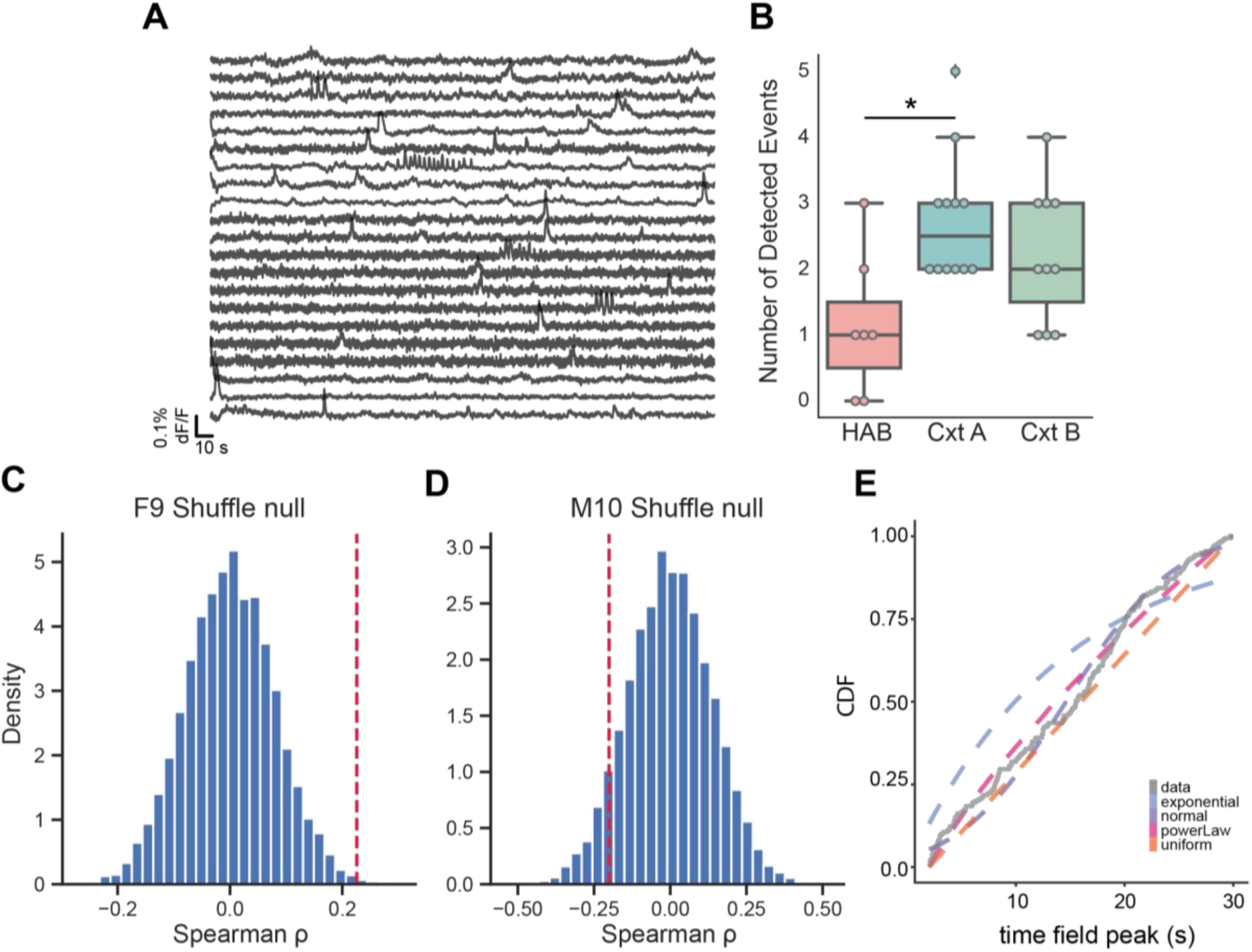
: Characterization of detected astrocytic calcium events during habituation to context A. **(A)** Example dF/F (%) traces sampled from the population of animals during habituation. **(B)** Comparison of number of detected events across conditions (HAB, Cxt A, Cxt B). Dots are individual animals. Significant differences between event numbers were detected (Kruskal-Wallis ANOVA p=0.020). Mann-Whitney pairwise comparisons found differences between HAB and Cxt A (p=0.022), no difference between HAB and Cxt B (p=0.089), and no difference between Cxt A and Cxt B (p= 0.458). [Hab: n=7, CxtA: n=11, CxtB: n=10] **(C-D)** Spearman’s rho shuffle null controls for F9 and M10, the only animals with at least two events. F9 displayed a significant correlation between events (p=0.005). M10 did not display between event correlation (p=0.149). (E) ECDF of data from F9 and M10 plotted against theoretical best fitting models. Uniform and Power-law were best fit distributions (WAIC [orange]=1192.2 and WAIC [pink]=1189.1 respectively).

## References.

1. O’Keefe, J. & Dostrovsky, J. The hippocampus as a spatial map. Preliminary evidence from unit activity in the freely-moving rat. Brain Res 34, 171–175 (1971).

2. O’Keefe, J. Place units in the hippocampus of the freely moving rat. Exp Neurol 51, 78–109 (1976).

3. O’Keefe, J. & Speakman, A. Single unit activity in the rat hippocampus during a spatial memory task. Exp Brain Res 68, 1–27 (1987).

4. McNaughton, B. L., Battaglia, F. P., Jensen, O., Moser, E. I. & Moser, M.-B. Path integration and the neural basis of the ‘cognitive map’. Nat Rev Neurosci 7, 663–678 (2006).

5. Eichenbaum, H. Time cells in the hippocampus: a new dimension for mapping memories. Nat Rev Neurosci 15, 732–744 (2014).

6. Mau, W., et al. The Same Hippocampal CA1 Population Simultaneously Codes Temporal Information over Multiple Timescales. Curr Biol 28, 1499–1508.e4 (2018).

7. Curreli, S., Bonato, J., Romanzi, S., Panzeri, S. & Fellin, T. Complementary encoding of spatial information in hippocampal astrocytes. PLOS Biology 20, e3001530 (2022).

8. Doron, A., et al. Hippocampal astrocytes encode reward location. Nature 609, 772–778 (2022).

9. Polykretis, I. & Michmizos, K. P. The role of astrocytes in place cell formation: A computational modeling study. J Comput Neurosci 50, 505–518 (2022).

10. Adamsky, A., et al. Astrocytic Activation Generates De Novo Neuronal Potentiation and Memory Enhancement. Cell 174, 59–71.e14 (2018).

11. Li, Y., et al. Activation of astrocytes in hippocampus decreases fear memory through adenosine A1 receptors. Elife 9, e57155 (2020).

12. Kol, A., et al. Astrocytes contribute to remote memory formation by modulating hippocampal–cortical communication during learning. Nat Neurosci 23, 1229–1239 (2020).

13. Williamson, M. R., et al. Learning-associated astrocyte ensembles regulate memory recall. Nature 637, 478–486 (2025).

14. Dewa, K., et al. The astrocytic ensemble acts as a multiday trace to stabilize memory. Nature 648, 146–156 (2025).

15. Sheintuch, L., et al. Tracking the Same Neurons across Multiple Days in Ca2+ Imaging Data. Cell Rep 21, 1102–1115 (2017).

16. Suthard, R. L., et al. Basolateral Amygdala Astrocytes Are Engaged by the Acquisition and Expression of a Contextual Fear Memory. J Neurosci 43, 4997–5013 (2023).

17. Suthard, R. L., et al. Engram reactivation mimics cellular signatures of fear. Cell Rep 43, 113850 (2024).

18. Qin, H., et al. Monitoring Astrocytic Ca2+ Activity in Freely Behaving Mice. Front Cell Neurosci 14, 603095 (2020).

19. Paukert, M., et al. Norepinephrine controls astroglial responsiveness to local circuit activity. Neuron 82, 1263–1270 (2014).

20. Srinivasan, R., et al. Ca2+ signaling in astrocytes from Ip3r2−/− mice in brain slices and during startle responses in vivo. Nat Neurosci 18, 708–717 (2015).

21. Cao, R., Bladon, J. H., Charczynski, S. J., Hasselmo, M. E. & Howard, M. W. Internally generated time in the rodent hippocampus is logarithmically compressed. eLife 11, e75353 (2022).

22. Watanabe, S. Asymptotic Equivalence of Bayes Cross Validation and Widely Applicable Information Criterion in Singular Learning Theory.

23. Buzsáki, G. & Mizuseki, K. The log-dynamic brain: how skewed distributions affect network operations. Nat Rev Neurosci 15, 264–278 (2014).

24. MacDonald, C. J., Lepage, K. Q., Eden, U. T. & Eichenbaum, H. Hippocampal ‘time cells’ bridge the gap in memory for discontiguous events. Neuron 71, 737–749 (2011).

25. Cruzado, N. A., Tiganj, Z., Brincat, S. L., Miller, E. K. & Howard, M. W. Conjunctive representation of what and when in monkey hippocampus and lateral prefrontal cortex during an associative memory task. Hippocampus 30, 1332–1346 (2020).

26. Lee, E. K., et al. Non-linear dimensionality reduction on extracellular waveforms reveals cell type diversity in premotor cortex. eLife 10, e67490 (2021).

27. Lee, K., Carr, N., Perliss, A. & Chandrasekaran, C. WaveMAP for identifying putative cell types from *in vivo* electrophysiology. STAR Protocols 4, 102320 (2023).

28. Gomez, J. A., et al. Ventral tegmental area astrocytes orchestrate avoidance and approach behavior. Nat Commun 10, 1455 (2019).

29. Mu, Y., et al. Glia Accumulate Evidence that Actions Are Futile and Suppress Unsuccessful Behavior. Cell 178, 27–43.e19 (2019).

30. Cho, W.-H., et al. Hippocampal astrocytes modulate anxiety-like behavior. Nat Commun 13, 6536 (2022).

31. Noh, K., et al. Cortical astrocytes modulate dominance behavior in male mice by regulating synaptic excitatory and inhibitory balance. Nat Neurosci 26, 1541–1554 (2023).

32. Keiser, A. A., et al. Sex Differences in Context Fear Generalization and Recruitment of Hippocampus and Amygdala during Retrieval. Neuropsychopharmacology 42, 397–407 (2017).

33. Martin-Fernandez, M., et al. Synapse-specific astrocyte gating of amygdala-related behavior. Nat Neurosci 20, 1540–1548 (2017).

34. Ahmed, M. S., et al. Hippocampal Network Reorganization Underlies the Formation of a Temporal Association Memory. Neuron 107, 283–291.e6 (2020).

35. Lefton, K. B. et al. Norepinephrine Signals Through Astrocytes To Modulate Synapses. bioRxiv 2024.05.21.595135 (2024) doi:10.1101/2024.05.21.595135.

36. Cahill, M. K., et al. Network-level encoding of local neurotransmitters in cortical astrocytes. Nature 629, 146–153 (2024).

37. Modi, M. N., Dhawale, A. K. & Bhalla, U. S. CA1 cell activity sequences emerge after reorganization of network correlation structure during associative learning. eLife 3, e01982 (2014).

38. Zhang, L., et al. Dynamics of a hippocampal neuronal ensemble encoding trace fear memory revealed by in vivo Ca2+ imaging. PLOS ONE 14, e0219152 (2019).

39. Mount, R. A., et al. Distinct neuronal populations contribute to trace conditioning and extinction learning in the hippocampal CA1. Elife 10, e56491 (2021).

40. Shikano, Y., Ikegaya, Y. & Sasaki, T. Minute-encoding neurons in hippocampal-striatal circuits. Current Biology 31, 1438–1449.e6 (2021).

41. Giovannucci, A., et al. CaImAn an open source tool for scalable calcium imaging data analysis. eLife 8, e38173 (2019).

42. Wang, Y., et al. Accurate quantification of astrocyte and neurotransmitter fluorescence dynamics for single-cell and population-level physiology. Nat Neurosci 22, 1936–1944 (2019).

43. Rupprecht, P., et al. Centripetal integration of past events in hippocampal astrocytes regulated by locus coeruleus. Nat Neurosci 27, 927–939 (2024).

44. Reitman, M. E., et al. Norepinephrine links astrocytic activity to regulation of cortical state. Nat Neurosci 26, 579–593 (2023).

45. Chen, A. B., et al. Norepinephrine changes behavioral state through astroglial purinergic signaling. Science 388, 769–775 (2025).

46. Murphy-Royal, C., Ching, S. & Papouin, T. A conceptual framework for astrocyte function. Nat Neurosci 26, 1848–1856 (2023).

47. Lines, J. Astrocytes integrate time and space. Neural Regeneration Research 20, 467 (2025).

48. Maren, S., De Oca, B. & Fanselow, M. S. Sex differences in hippocampal long-term potentiation (LTP) and Pavlovian fear conditioning in rats: positive correlation between LTP and contextual learning. Brain Res 661, 25–34 (1994).

49. Gresack, J. E., Schafe, G. E., Orr, P. T. & Frick, K. M. Sex differences in contextual fear conditioning are associated with differential ventral hippocampal extracellular signal-regulated kinase activation. Neuroscience 159, 451–467 (2009).

50. Malik, R., Li, Y., Schamiloglu, S. & Sohal, V. S. Top-down control of hippocampal signal-to-noise by prefrontal long-range inhibition. Cell 185, 1602–1617.e17 (2022).

51. Etter, G., van der Veldt, S., Choi, J. & Williams, S. Optogenetic frequency scrambling of hippocampal theta oscillations dissociates working memory retrieval from hippocampal spatiotemporal codes. Nat Commun 14, 410 (2023).

52. Stringer, C., Wang, T., Michaelos, M. & Pachitariu, M. Cellpose: a generalist algorithm for cellular segmentation. Nat Methods 18, 100–106 (2021).

53. Jean-Richard-dit-Bressel, P., Clifford, C. W. G. & McNally, G. P. Analyzing Event-Related Transients: Confidence Intervals, Permutation Tests, and Consecutive Thresholds. Front. Mol. Neurosci. 13, (2020).

54. Mathis, A., et al. DeepLabCut: markerless pose estimation of user-defined body parts with deep learning. Nat Neurosci 21, 1281–1289 (2018).

55. Pedregosa, F., et al. Scikit-learn: Machine Learning in Python. Journal of Machine Learning Research 12, 2825–2830 (2011).

